# Coat protein of TCV suppresses RNA decay via ubiquitination and autophagy pathways to facilitate viral infection

**DOI:** 10.1101/2024.09.05.611546

**Authors:** Kunxin Wu, Qiuxian Xie, Xueting Liu, Yan Fu, Shuxia Li, Xiaoling Yu, Wenbin Li, Pingjuan Zhao, Yanli Ren, Mengbin Ruan, Xiuchun Zhang

**Affiliations:** National Key Laboratory for Tropical Crop Breeding, Institute of Tropical Bioscience and Biotechnology, Chinese Academy of Tropical Agricultural Sciences & Key Laboratory for Biology and Genetic Resources of Tropical Crops of Hainan Province, Hainan Institute for Tropical Agriculture Resources, Haikou, 571101, China; College of Biological Sciences and Technology, Yili Normal University, Yining 835000, China; Sanya Research Institute, Chinese Academy of Tropical Agricultural Sciences, Sanya, 572025, China

**Keywords:** *Turnip crinkle virus* (TCV), viral suppressor of RNA decay, antiviral, decapping 1 (Dcp1), protein degradation

## Abstract

**Highlight statement:** The findings highlight that TCV manipulates 26S proteasome and autophagy pathways to obstruct antiviral RNA decay defenses and ultimately enhance its ability to infect host cells.

RNA decay is a pervasive process in eukaryotic cells. Viruses utilize the host cell’s intracellular machinery to gain access to essential molecules and subcellular structures required for infection during the pathogenesis process. The study demonstrates that turnip crinkle virus (TCV) infection enhances the expression of Arabidopsis Dcp1 (AtDcp1), which negatively regulates the accumulation of TCV RNA, indicating its involvement in antiviral defense. Nevertheless, TCV circumvents the antiviral defense based on RNA decay, as indicated by the capsid protein (CP) of TCV stabilizing the known nonsense-mediated RNA decay targeted transcripts. In vivo, CP physically interacts with AtDcp1, promoting AtDcp1 degradation via ubiquitination and autophagy pathways. This is evidenced by the observation that the degradation is inhibited by both 26S proteasome and autophagy inhibitors. Furthermore, CP elevates the polyubiquitination of Dcp1-Flag and the quantity of pre-autophagosome or autophagosome structures. These data indicate that CP suppresses RNA decay by interacting with AtDcp1 and mediating its degradation through the 26S proteasome and autophagy pathways, effectively subduing antiviral RNA decay. This study uncovers a previously unidentified virulence strategy in the ongoing conflict between plants and viruses.

## INTRODUCTION

Plants are frequently subjected to a variety of biotic and abiotic stresses, including viral infections. However, throughout their extensive co-evolution with viruses, plant cells have evolved intricate and distinct defense mechanisms, such as RNA silencing and RNA decay, directly target viral nucleic acids (Zhang et al., 2017). Furthermore, a number of regulatory mechanisms oversee protein homeostasis, including the ubiquitin-proteasome-degradation system (UPS) and autophagy, which play a role in combating viral invasion (Wu et al., 2023a; Sun et al, 2023). RNA decay is a common process presents in eukaryotic cellsand plays a pivotal role in determining the fate of cellular RNAs. In order to enhance their coding potential, a considerable number of RNA viruses have evolved to possess multi-cistronic genomes. Therefore, viruses with a lengthy 3′ UTR (e.g. >3000 nt) are highly susceptible to nonsense-mediated RNA decay (NMD) (Balistreri *et al*., 2017; Hogg, 2016). Plants deficient in RNA decay-related factors are more susceptible to viral infections, which highlighted the crucial role of NMD as a cellular antiviral mechanism (Li and Wang, 2018; 2019; May and Simon, 2021; Wu et al., 2023a; 2023b). The core factor in both nonstop decay (NSD) and no-go decay (NGD) processes effectively recognize the G2A6 motif present in the P3 coding region of three different potyviruses, resulting in the cleavage of viral RNA (Ge et al., 2023). This demonstrates the antiviral defense of RNA decay in plant.

Viruses are obligatory intracellular parasites that cannot reproduce outside of host cells. In the ongoing molecular arms race between host cells and viruses, the latter have developed various strategies to evade or counteract the natural antiviral defense mechanism of the host (Guo *et al*., 2018; Li and Wang, 2019; Popp *et al*., 2020; May et al., 2021; Wu et al., 2023a). These strategies may differ among virus types and may be executed via cis-acting or trans-acting methods. Cis-acting elements are primarily composed of viral RNA sequences or structures that evade the host RNA decay process (May et al., 2018). In contrast, trans-acting molecules include viral proteins that target or hijack RNA decay signaling components, thereby enhancing viral infection (Zhang et al., 2020; May et al., 2020; Wu et al., 2023b).

Accumulating evidence indicates that the ubiquitin-proteasome-degradation system (UPS) and autophagy play a dual role in combating and promoting viral infections (Wu et al., 2023a; Yang et al., 2022a). These processes help to maintain plant health by balancing viral infection with host survival in the intense struggle between host plants and viral pathogens. Plant viruses have also evolved to manipulate host UPS and autophagy processes in their favor. For instance, viruses can impede the degradation of their own proteins or accelerate the degradation of RNA silencing components to facilitate their own spread and infection (Cheng and Wang, 2016; Derrien et al., 2012; Li et al., 2017).

*Turnip crinkle virus* (TCV) belongs to the *Betacarmovirus* genus of the Tombusviridae family. It is reported that the TCV genomic RNA is resistant to RNA decay as a result of a read-through structure following the termination codon in the p28 mRNA and an unstructured region of 51 nucleotides at the beginning of the TCV 3’ UTR (May *et al*. 2018). The 38 kDa capsid protein (CP) serves multiple functions. The CP encoded by TCV is not only a determinant of viral symptoms but also a suppressor of RNA silencing. Wu et al. (2023b) discovered that it inhibits RNA degradation by binding with Arabidopsis XRN4, resulting in enhanced viral infection. However, the mechanisms by which CP counters plant antiviral RNA decay defense to promote TCV infection remain unclear.

This study reports that TCV infection increases the fluorescence intensity of AtDcp1, which negatively regulates TCV RNA accumulation. Additionally, CP protein interacts with AtDcp1. Though the interaction did not impair the formation of the Dcp1/Dcp2 decapping complexes, the present of CP causes a decrease in the quantity of AtDcp1 granules. Furthermore, evidences indicate that CP mediates AtDcp1 degradation via both 26S proteasome and autophagy pathways resulting in stabilization of the known NMD targets and facilitating viral infection. These findings demonstrate that TCV manipulate 26S proteasome and autophagy pathways to obstruct antiviral RNA decay defense and ultimately enhance its ability to infect host cells.

## RESULTS

### PBs and exosome complexes are altered in plants infected with TCV

The process of RNA decay involves a series of coordinated molecular events (Furuichi et al., 1977; Hsu and Stevens, 1993). Decapping is generally a prerequisite for the degradation of most RNA by the 5’-3’ XRN exoribonuclease exosome complex (Tsuzuki et al., 2017; Zhang and Guo, 2017). In *Arabidopsis thaliana* (Arabidopsis), the decapping process is carried out by the nudix hydrolase DECAPPING2 (Dcp2) and its associated cofactors Dcp1 and VARICOSE (VCS) (Xu et al., 2006). Several proteins, including Dcp5 and the SM-like (LSM) 1-7 complex, are involved in decapping activation and processing body (PB) assembly (Xu and Chua, 2009; Perea-Resa et al., 2012). To investigate the role of decapping in defending against TCV, we conducted a transient expression assay in *N. benthamiana* cells using agroinfiltration. The subcellular localization of RNA granules in *N. benthamiana* epidermal cells was initially examined by transiently expressing AtDcp1 and AtRRP41 fused with fluorescent reporters (GFP) (refer to Fig. 1a). AtDcp1 and AtRRP41 are marker genes for PBs and potentially marked RNA-exosome core complexes, respectively. As shown in Figure 1b, the markers showed a cytoplasmic distribution with varying degrees of condensation into droplet-like foci under mock conditions (top panel). The AtDcp1 and AtRRP41 fusion were primarily assembled in the foci. These localization patterns were similar to those previously described (Xu *et al*., 2006; Motomura *et al*., 2015). It is noteworthy that the accumulation of enhanced GFP was observed to be enhanced in plants infected with TCV, as evidenced by both the Dcp1-GFP and RRP41-GFP fusions. In contrast, the expression of free GFP by pG1300 was found to be unaltered. Statistical analyses were conducted by measuring the fluorescence intensity in the designated regions of interest, with three to five images captured for each treatment and over 15 measurements taken for each image. The data demonstrated that TCV infection markedly elevated the fluorescence intensity of AtDcp1 and AtRRP41 (Figure 1c), a finding that aligns with previous reports indicating that TMV and CaMV infection enhances the fluorescence intensity in regions of interest with PBs and RRP41 (Conti et al., 2017; Hoffmann et al., 2022). In conclusion, it can be hypothesized that the alteration of different components of RNA granules may be part of a plant defense response, induced to degrade and reduce the accumulation of viral RNAs.

**FIGURE 1.**
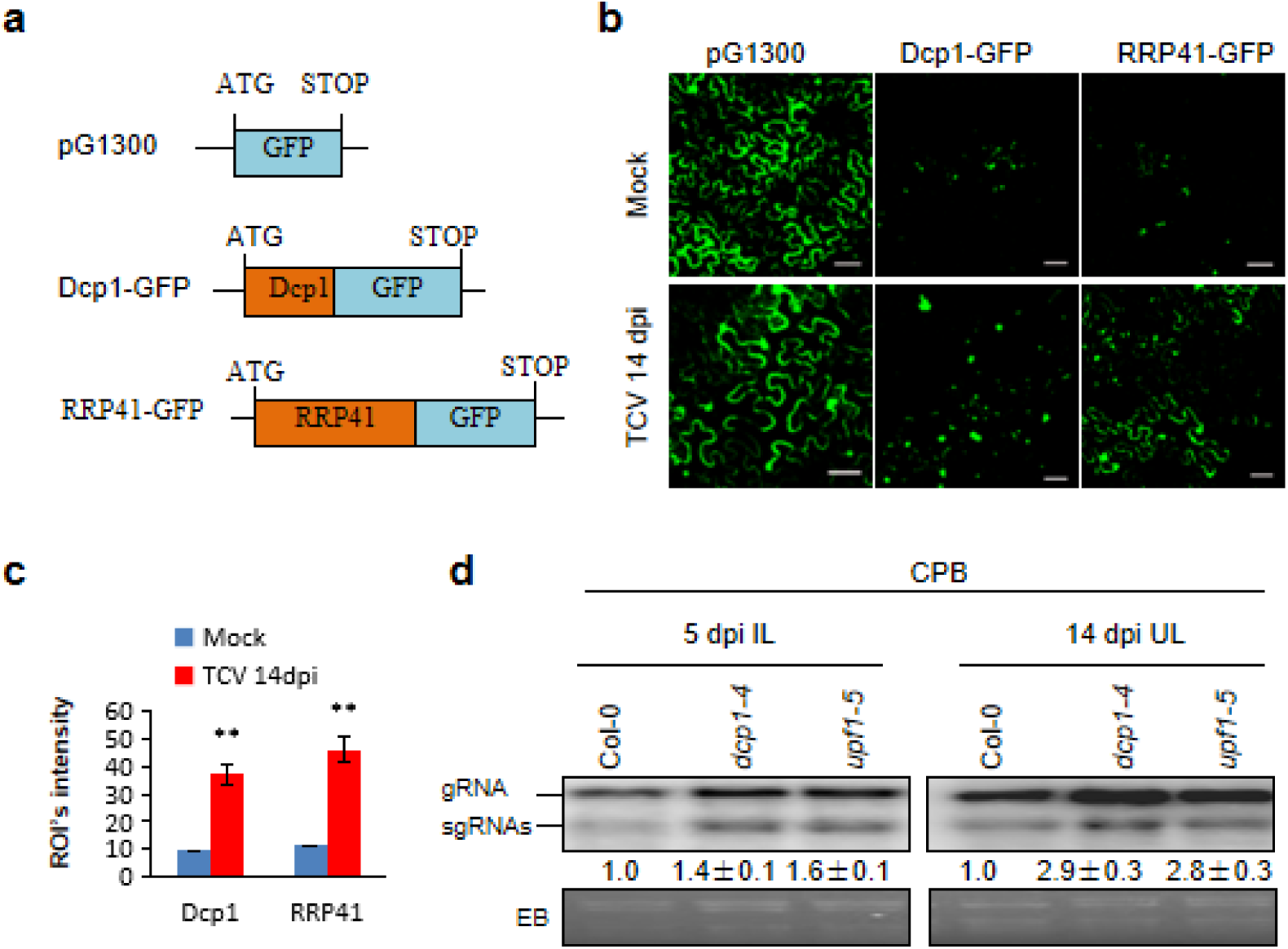
RNA decay was involved in anti-TCV defense in plants. (a) Schematic representation of free GFP (pG1300), AtDcp1 or AtRRP41 fusion with GFP. (b) Confocal section images showing subcellular localization of the indicated fluorescent fusion proteins in *N. benthamiana* leaves of mock or TCV-infected plants at 14 days past agroinfiltration (dpa). Bar, 50 μm. (c) Fluorescence intensity of several regions of interest (ROIs intensity level) in AtDcp1 expressing cells. Statistical analyses were performed with three to five images for each treatment and more than 15 measurements for each picture. Values presented represent the mean ± standard deviation (SD). Double asterisks indicate a highly significant difference (p<0.01) between TCV-infected plants at 14 dpa and mock (Student’s *t* test). (d) Detection of genomic RNA (gRNA) and subgenomic RNA (sgRNAs) of CPB by Northern blot hybridization in inoculated leaves (ILs) and upper non-inoculated leaves (ULs) from Col-0, *dcp1-4* and *upf1-5* mutant plants infected with the infectious clone of TCV mutant CPB, which contained single aa change at aa residue 130 (R130T) of TCV CP, respectively, and kept at 18 ℃. The probe was DIG-labeled DNA, which is a 667-nt fragment of TCV. The relative amount of CPB gRNA and sgRNA is indicated. Each experiment was repeated at least three times. EB: ethidium bromide-stained Northern gel. See Wu *et al*. (2021) for a detailed procedure.

### AtDcp1 negatively regulates TCV RNA accumulation

To further investigate the potential contribution of AtDcp1 to anti-TCV defense, we obtained the Arabidopsis *dcp1-4* knockout mutant (Li & Wang, 2018) and *upf1-5* mutant plants, in which AtUpf1 mRNA levels were severely reduced (Lukhovitskaya & Ryabova, 2019). The mutants were confirmed by genotyping. It is noteworthy that CP is a potent viral suppressor of RNA silencing (VSR), which shuts down the main pathway (Qu et al., 2003). Consequently, only TCV mutants with significantly weaker VSR, such as CPB and CPC, can reveal the anti-TCV functions of genes such as DCL4 and DCL2 (Qu et al., 2008; Zhang et al., 2012). Therefore, we inoculated plants with *in vitro* transcribed infectious RNA of TCV mutant CPB. The CPB virus RNA levels in Arabidopsis were much lower at 26 °C than at 18 °C due to reduced CPB VSR activity combined with increased host RNA silencing activity (Zhang et al., 2012). As a result, the inoculated plants were maintained at 18℃. Northern blot hybridization revealed that the accumulation levels of CPB RNAs increased at five days post inoculation (dpi) in inoculated leaves (ILs) and 14 dpi in upper non-inoculated leaves (ULs) of both *dcp1-4* and *upf1-5* mutant plants in comparison to their wild-type counterparts. (Fig. 1d). These findings indicate that AtDcp1, a co-factor of the decapping complex, negatively impacts the accumulation of TCV RNA as AtUpf1 which is an essential factor for nonsense-mediated mRNA decay (NMD).

### CP Interacts with *At*Dcp1

The aforementioned outcomes indicated that AtDcp1 has an inhibitory effect on the accumulation of TCV RNA. Nevertheless, TCV can still infect Arabidopsis, indicating the presence of an antidote defense strategy employed by TCV. It was postulated that viral protein(s) may be implicated in this counterdefense. Given that CP is a potent suppressor of RNA silencing, we sought to ascertain whether it might impede the deleterious effects of AtDcp1 through a physical interaction. The aim of this study was to investigate the physical interaction between CP and AtDcp1 using yeast two-hybrid assays (Y2H). Robust growth was observed on the QDO medium containing 7.5 mM 3-AT when CP, CPB, or CPC were linked to the AD vector and AtDcp1 was linked to the BD vector, as illustrated in Fig. 2a. It is noteworthy that although the BD-Dcp1 construct exhibited autoactivation, the use of 7.5 mM 3-AT in the selection medium effectively prevented this phenomenon. The lack of interaction between TCV P28 and AtDcp1 was demonstrated by the inability of the AD-P28/BD-Dcp1 combination to grow on selective media containing 7.5 mM 3-AT, which also served as a negative control. These findings provide evidence in support of the hypothesis that CP, CPB, and CPC interact with AtDcp1 in yeast.

**FIGURE 2.**
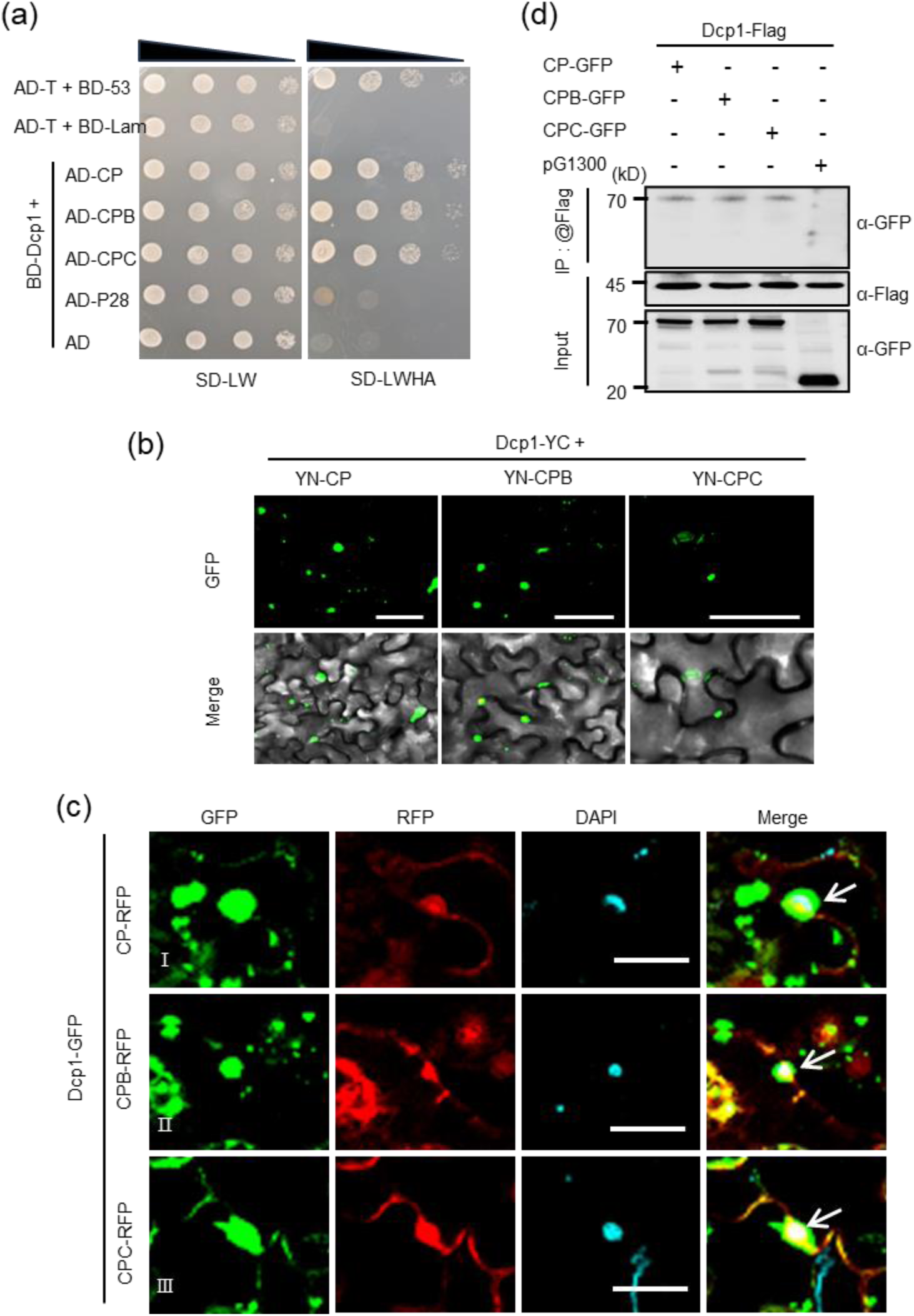
TCV CP interacted with Arabidopsis Dcp1. (a) Yeast two-hybrid assay (Y2H) for protein–protein interactions between BD-Dcp1 and AD-CP or its mutant AD-CPB or AD-CPC on the quadruple dropout (QDO) medium SD−LWHA containing 7.5 mM 3-amino1,2,4-triazole (3-AT). The positive and negative controls are the yeast co-transformants with pGAD-T plus pGBK-53 and pGAD-T plus pGBK-Lam, respectively. AD, GAL4 activation domain; BD, GAL4 DNA binding domain; SD-LWHA, synthetic defined (SD) yeast minimal medium lacking Leu, Trp, His, and adenine hemisulfate. (b) Bimolecular fluorescence complementation (BiFC) assay for protein–protein interactions between TCV CP (I) or its mutant CPB (II) or CPC (III) and AtDcp1. TCV CP, CPB, and CPC were fused with the N-terminal, while AtDcp1 was fused with the C-terminal half of YFP, designated as YN-CP, YN-CPB, YN-CPC and Dcp1-YC, respectively, and transiently expressed in *N. benthamiana* leaves. (c) Confocal micrographs of *N. benthamiana* co-expressing (CP-RFP) (Ⅰ), CPB (CPB-RFP) (Ⅱ) or CPC (CPC-RFP) (Ⅲ) with Dcp1-GFP at 48 hpa. Bar, 50 μm. White arrows represent nuclear interaction body, respectively. (d) Co-immunoprecipitation (Co-IP) detection of interaction between AtDcp1 and viral protein CP or its mutant CPB, CPC in *Nicotiana benthamiana* leaves. Dcp1-Flag and CP-GFP, CPB-GFP, CPC-GFP or pG1300 (negative control) were transiently co-expressed in wildtype *N. benthamiana* leaves. After 2 days of expression, proteins were extracted and immunoprecipitated with Flag-trapped agarose. The input proteins were analyzed using anti-GFP and anti-Flag antibodies, while the immunoprecipitated (IP) proteins were analyzed using anti-GFP antibody.

To ascertain whether CP, CPB, or CPC interacted with AtDcp1 in planta, then bimolecular fluorescence complementation (BiFC) assay was conducted. As illustrated in Figs 2b & S1a, a strong interaction between CP, CPB, or CPC and AtDcp1 was observed in both nucleus and cytoplasm. AtDcp2 consistently interacted with AtDcp1, functioning as a positive control, and generating vivid granules as previously described by Xu et al. (2006), while no interaction was observed when performing experiments with corresponding control constructs (Dcp1-YC with YN-P28) (Fig. S1b). Subcellular localization assays have also indicated that both CP, CPB, and CPC colocalize with AtDcp1 (Figs. 2c and S2). The interaction between CP, CPB, or CPC and AtDcp1 was further validated by a immunoprecipitation (Co-IP) assay. The Flag-tagged AtDcp1 protein (Dcp1-Flag) was transiently co-expressed with CP, CPB, CPC-GFP, or with pG1300 with free GFP in *N. benthamiana* plants. As illustrated in Figure 2d, the Co-IP analyses demonstrated that Dcp1-Flag Co-IP with both CP-GFP, CPB-GFP, and CPC-GFP, but not with free GFP (Fig. 2d), thereby confirming the interaction between CP, CPB, or CPC and AtDcp1 *in vivo*. In conclusion, these results demonstrate that Dcp1 interacts with CP, CPB, and CPC in vivo. Furthermore, despite the diminished silencing suppressor activity observed in CPB and CPC mutants, their interaction with AtDcp1 remained unaltered. Consequently, our findings indicate that the interaction between CP and AtDcp1 occurs independently of its VSR activity.

### CP did not disrupt the interaction between AtDcp1 and AtDcp2

As CP interacts with AtDcp1, we first examined whether it negatively affects the formation of the Dcp1/Dcp2 decapping complexes which are RNA decay sites. We analyzed the effects of CP on the Dcp1-Dcp2 interaction using a competitive BiFC assay. For the competitive BiFC assay, we co-expressed YN-Dcp2 and Dcp1-YC with CP-RFP or an empty vector pR1300 with free RFP (as a control) in *N. benthamiana* plants. Interaction signals between YN-Dcp2 and Dcp1-YC were observed at two days post-agroinfiltration (dpa), resulting in the reconstitution of YFP signals in both scenarios (Fig. 3a). To eliminate the potential impact of the RFP tag, we co-infiltrated YN-Dcp2 and Dcp1-YC with CP fused with an HA tag or an empty vector, along with Dcp1-RFP. The reconstituted YFP signals were colocalized with Dcp1-RFP (Fig. 3b). The CP expression was confirmed via an immunoblotting assay using an HA antibody, as illustrated in Figure 3C. As anticipated, the addition of CP markedly enhanced the formation of AtDcp1/AtDcp2 granules in the cytoplasm, in comparison with the vector control (Fig. 3d). This suggests that CP enhances the interaction between AtDcp1 and AtDcp2, potentially due to its VSR activity. These results indicate that CP does not disrupt the interaction between AtDcp1 and AtDcp2, which is in contrast to the effect observed with TuMV VPg. TuMV VPg has been shown to disturb the interaction between NbDcp1 and NbDcp2 by shifting NbDcp2 from cytoplasmic to nuclear targeting (Li et al., 2018).

**FIGURE 3.**
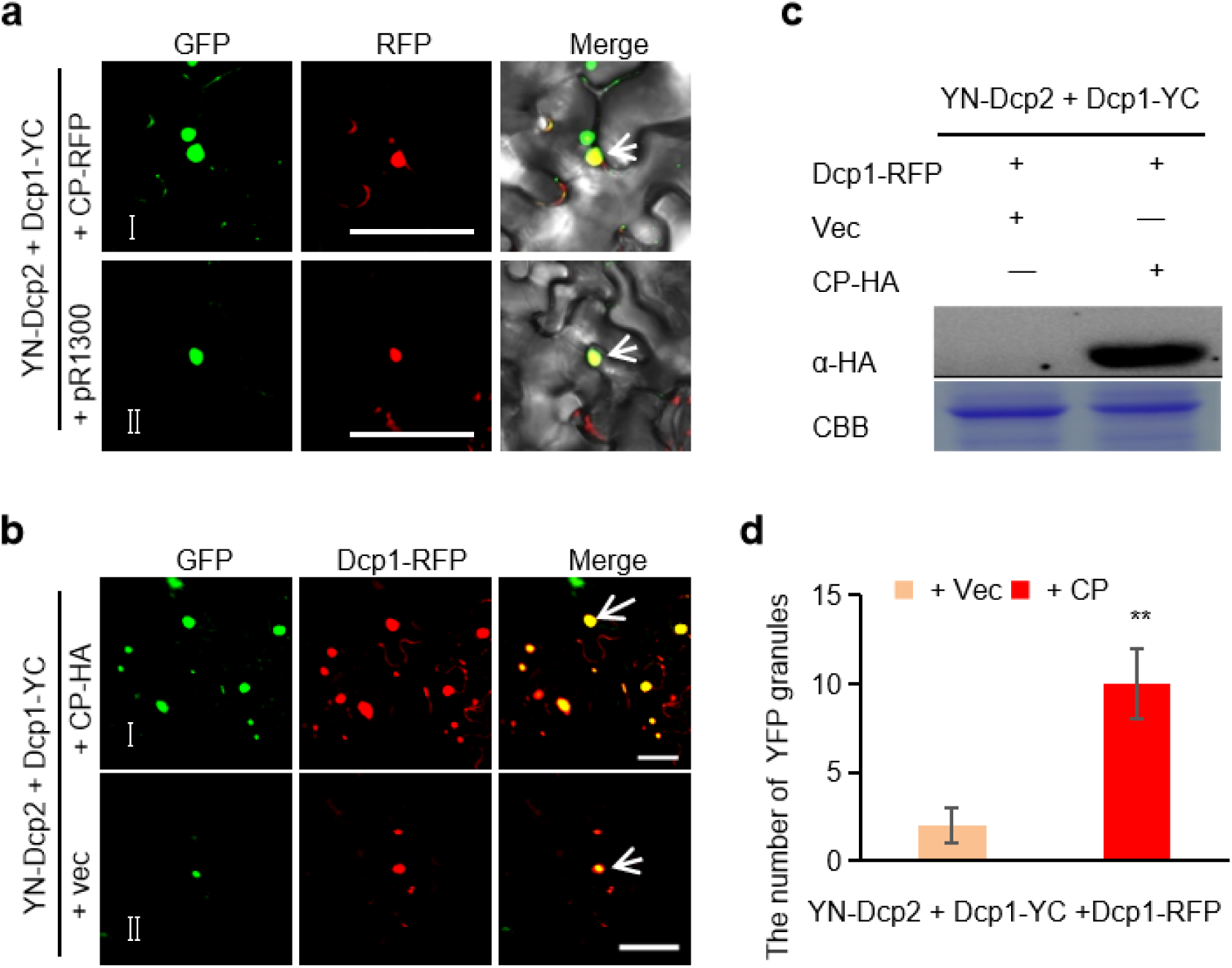
TCV CP failed to disrupt the interaction between AtDcp1 and AtDcp2. (a) Colocalization of CP-RFP or free RFP with reconstituted YFP from the BiFC assays for YN-Dcp2 and Dcp1-YC interaction. BiFC and colocalization assays were carried out in wild-type *N. benthamiana* leaf cells. Confocal microscopy was carried out at 2 dpa. White arrows indicated the colocalization of RFP and reconstituted YFP from BiFC assays for YN-Dcp2 and Dcp1-YC interaction. Bars, 50 μm. (b) Colocalization of Dcp1-RFP with reconstituted YFP from the BiFC assays for YN-Dcp2 and Dcp1-YC interaction in the presence of an empty vector (+ vec) or CP (+ CP-HA). BiFC and colocalization assays were carried out in wild-type *N. benthamiana* leaf cells. Confocal microscopy was carried out at 2 dpa. White arrows indicated the colocalization of Dcp1-RFP and reconstituted YFP from BiFC assays for YN-Dcp2 and Dcp1-YC interaction. Bars, 50 μm. (c) Western blot analysis using anti-HA antibody to confirm the CP expression in the infiltrated *N. benthamiana* leaf. Coomassie brilliant blue staining (CBB) of rubisco large subunit was used as a loading control. (d) The number of YFP granules reconstituted from YN-Dcp2 and Dcp1-YC interaction (per 10 cells), which colocalized with Dcp1-RFP, when they were co-infiltrated with vec (+vec) or CP (+CP). Independent infiltration experiments were repeated three times and 30 cells in total were used to quantify. Values represent the mean number of the YFP granules (per 10 cells) ± SD. Student’s t test was performed to compare differences, and double asterisks indicate a highly significant difference (P < 0.01).

### The expression of CP causes a decrease in the quantity of AtDcp1 granules and AtDcp1 protein accumulation independent of its VSR activity

It has been reported that the viral protein may impede the host’s antiviral system by breaking down host proteins, thereby facilitating infection during pathogenesis (Cheng et al., 2016; Li et al., 2017; Ji et al., 2021; Tong et al., 2021). The Dcp1/2 holoenzyme’s activity is meticulously monitored and precisely calibrated to align with the developmental and metabolic states of a cell, as the 5’-cap plays a pivotal role in the transcript lifecycle and the crucial step of decay (Vidya and Duchaine, 2022). In order to ascertain the mechanism by which TCV CP inhibits RNA decay, an initial examination was conducted to determine whether it affects the stability of AtDcp1. To assess the potential influence of CP on AtDcp1 stability, we conducted an experiment wherein pG1300 or Dcp1-GFP fusion proteins were co-expressed in the absence or presence of CP in the foliage of N. benthamiana. To eliminate the potential confounding effect of RNA silencing targeting the mRNA of all constructs, we co-infiltrated p19 with each assay. p19 is the viral suppressor of tomato bushy stunt virus (TBSV) that does not interfere with nonsense-mediated mRNA decay (NMD). Images captured at 4 dpa revealed no notable discrepancies in the dispersion of free GFP among tissues that were co-expressing with CP or not (Fig. 4a). The Dcp1-GFP fusion protein formed cytoplasmic granules when co-expressed in the presence or absence of CP. However, the co-expression of Dcp1-GFP with CP resulted in a notable reduction in the number of Dcp1-GFP granules, in comparison to plants expressing Dcp1-GFP with buffer (Fig. 4a and 4b). To ascertain whether the reduction in GFP fluorescence observed in plants co-expressing CP was attributable to a decline in Dcp1-GFP levels, we conducted Western blot analysis to confirm the accumulation of Dcp1-GFP at 4 dpa. The results of the analysis demonstrated that co-expression with CP resulted in a twofold decrease in AtDcp1 levels in comparison to that with buffer, as illustrated in Fig. 4c. These findings indicate that the overexpression of CP results in a reduction in AtDcp1 protein accumulation. Furthermore, the quantity of AtDcp1 generated in the presence of CP exhibited a gradual reduction over time (Fig. 4d). Similarly, in a cell-free degradation system, the degradation rate of AtDcp1 was significantly higher in the presence of GFP-CP than in the control GFP (Fig. 4e). Therefore, it can be concluded that CP protein promotes the degradation of AtDcp1.

**FIGURE 4.**
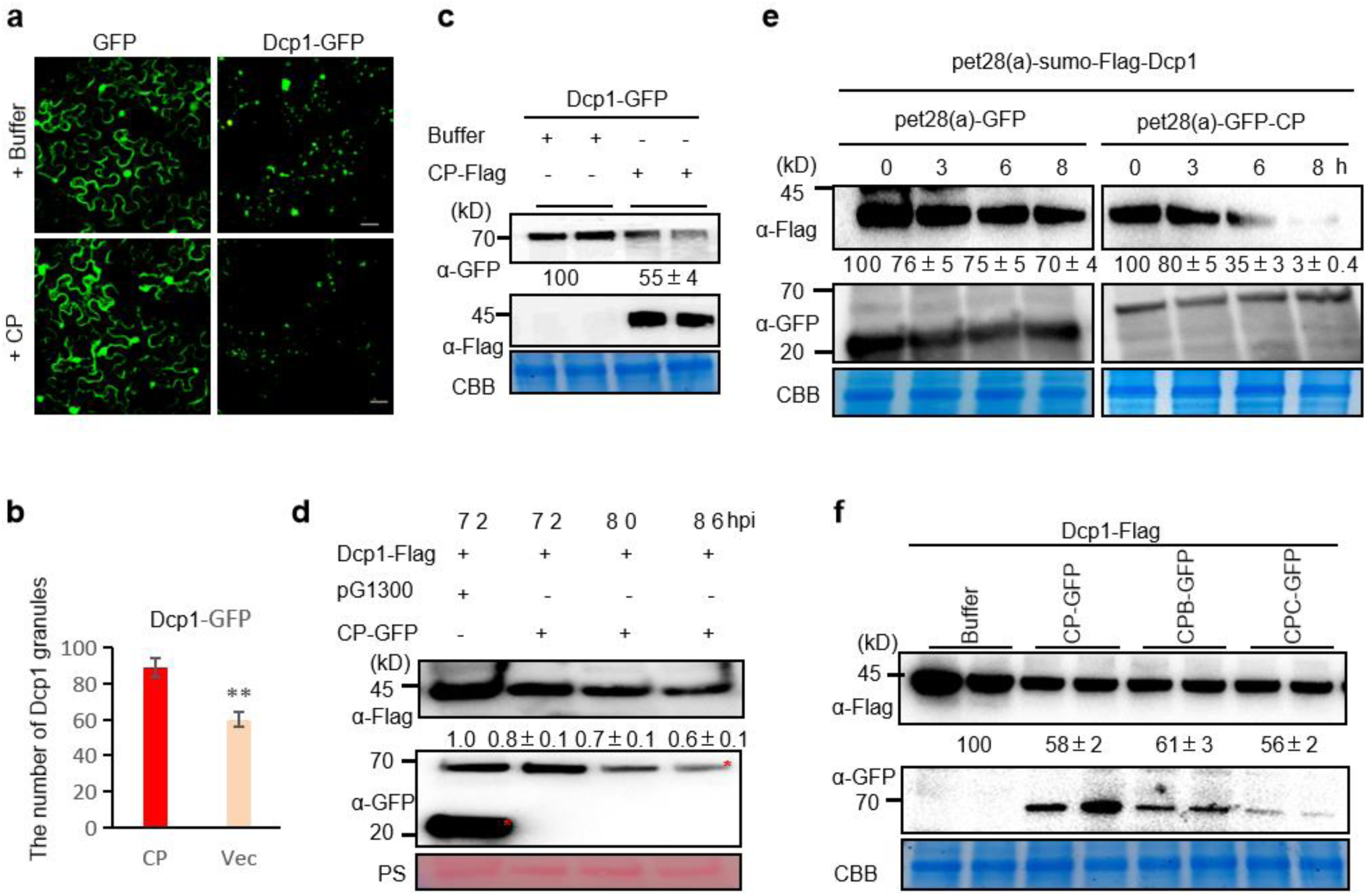
TCV CP-mediated degradation of AtDcp1 independent of its VSR activity. (a) Micrographs show cells expressing free GFP or Dcp1-GFP in conjunction with buffer (upper panel) or CP (lower panel). Infiltrated *N*. *benthamiana* leaves were examined at 4 dpa. Bars, 50 μm. (b) The average number of Dcp1-GFP spots per 3 cells. Infiltration experiments were repeated three times, and 30 cells were counted for the punctate spots. The average number was calculated using 3 cells as a unit. Values represent the mean spot ±standard deviation (SD) per 3 cells. Statistical analysis was performed using Student’s t-test (two-sided, * p < 0.05, ** p < 0.01). (c) Western blot of total protein extracts from (a) detected with GFP antibodies (top) or Flag antibodies (middle). (d) Time course degradation of AtDcp1 with or without CP *in vivo*. Western blot of total protein extracts from *N. benthamiana* leaves which co-infiltrated with Dcp1-Flag with CP-GFP, CP-GFP, CP-GFP or pG1300 with Flag antibodies (top) or GFP antibodies (middle). Samples were collected at the indicated times. Ponceau S (PS) staining served as a loading control. The asterisk symbol denotes the target band. (e) Time course degradation of AtDcp1 with or without CP *in vitro*. The crude protein extracts of pet28(a)-sumo-Flag-Dcp1 were incubated with same amount of pet28(a)-GFP or pet28(a)-GFP-CP crude protein extracts at 37 °C, and samples were collected at the indicated times for western blot using anti-Flag and anti-GFP. CBB served as a loading control.(f) Western blot of total protein extracts from *N. benthamiana* leaves which co-infiltrated with Dcp1-Flag with CP-GFP, CP-GFP, CP-GFP or pG1300 at 4 dpa. (c) - (f) the protein assays were done at least three independent times, and protein bands from two representative experiments were quantified using Image J software. The amount of protein detected in samples co-expressing with buffer or pG1300 was set at 100 and values under the blots represent the average level of protein. Ponceau S (PS) stainning of rubisco large subunit or CBB shows equal protein loading.

Given that the binding between CP and AtDcp1 is independent of its VSR activity, we sought to determine whether the reduction in AtDcp1 protein accumulation resulting from CP is also independent of its VSR activity. To eliminate any potential confounding effects of the GFP tag, it was substituted with the Flag tag on AtDcp1. Subsequently, *N. benthamiana* leaves were agroinfiltrated with Dcp1-Flag and buffer, CP-GFP, CPB-GFP, or CPC-GFP, in the presence of TBSV p19. Samples were harvested at 4 dpa. As illustrated in Figure 4f, the co-expression of CP, CPB, and CPC resulted in a comparable reduction in AtDcp1 levels relative to the buffer control. These findings indicate that the reduction in AtDcp1 protein accumulation resulting from CP is also independent of its VSR activity.

### CP mediates AtDcp1 degradation via both 26S proteasome and autophagy pathways

The UPS and autophagy pathways are conserved cellular degradation pathways that regulate a multitude of biological processes in plants. To determine whether CP-mediated AtDcp1 protein degradation occurs via the UPS or autophagy pathway, the sensitivity of CP or AtDcp1 to MG32 and 3-MA, two chemical inhibitors of the 26S proteasome and autophagy pathway, respectively, was evaluated. Leaves of *N. benthamiana* were agroinfiltrated with Dcp1-Flag or CP-GFP, along with TBSV p19, and then infiltrated with either DMSO (control) or MG132 (100 μM) or 3-MA (10 mM) after 60 hours. Leaf samples were subsequently collected after an additional 12-hour incubation period. Samples collected at 4 dpa demonstrate that treatment with MG132 resulted in a 20% increase in AtDcp1 expression in N. benthamiana leaf tissues, while treatment with 3-MA increased it by 8% (Fig. S3a).

Therefore, AtDcp1 turnover maybe link to both 26S proteasome and autophagy pathways. However, no discernible alteration in CP-GFP protein levels was discerned in samples treated with DMSO, MG132, or 3-MA (Fig. S3b). These results indicate that the inhibition of the 26S proteasome and autophagy may not affect the accumulation of CP when expressed alone. Subsequently, additional experiments were conducted to investigate the impact of MG132 on the accumulation of AtDcp1 and CP in *N. benthamiana* leaves when co-expressed. As illustrated in Fig. 5a, the AtDcp1 level exhibits a pronounced decline in the presence of CP, which is consistent with the observations depicted in Fig. 4. These observations corroborate the reduction of AtDcp1 induced by CP. However, the reduction of AtDcp1 mediated by CP is prevented by the addition of the 26S proteasome inhibitor MG132 (Fig. 5a, top panel). Furthermore, MG132 treatment did not affect CP that was co-expressed with AtDcp1 (Fig. 5a, middle panel), which is consistent with the results obtained from independently expressed CP (Fig. S3b). These findings suggest that the reduction of AtDcp1 by CP may occur via the 26S proteasome pathway.

**FIGURE 5.**
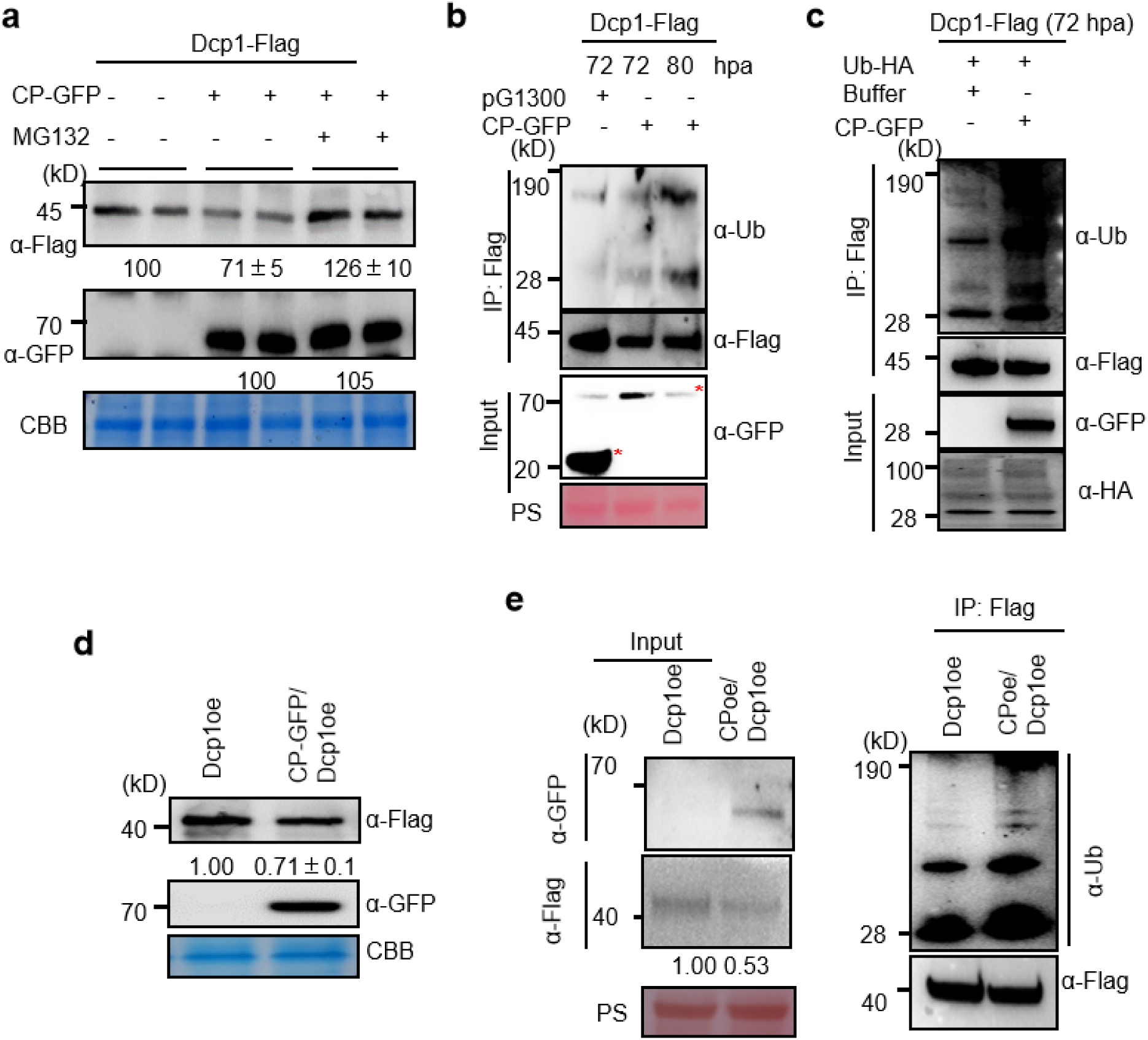
CP mediates AtDcp1 degradation via 26S proteasome pathway. (a) CP-triggered AtDcp1 degradation was blocked by proteasome inhibitor MG132. Dcp1-Flag was transiently expressed alone or with CP-GFP in *N. benthamiana* leaves. At 72 hpa, the infiltrated leaf areas were treated with MG132 or DMSO for an additional 12 h. Total protein was extracted, separated by SDS-PAGE, and detected with Flag antibodies or GFP antibodies. The relative amount of AtDcp1 or CP is indicated. The bottom panel shows equal protein loading as determined by CBB staining of rubisco. Each experiment was repeated at least three times. (b,c) CP enhances the process of ubiquitination of AtDcp1. Dcp1-Flag was co-expressed with pG1300 or CP in absent (b) or present of Ub-HA (c) in *N. benthamiana*. At 72 hpa, the infiltrated leaf areas were treated with DMSO or MG132 for an additional 12 or 18 hpa, and then the extracts were obtained for western blot using anti-GFP or anti-HA antibody and were precipitated with Protein A/G Flag antibody beads. Similar amounts of Dcp1-Flag precipitated by the anti-Flag antibody beads were used for analysis by immunoblotting using anti-Flag antibody or anti-Ubiquitin (Ub) respectively. PS was used as a loading control to monitor input protein amounts. The asterisk symbol denotes the target band. (d) Expression of Dcp1-Flag, CP-GFP in Arabidopsis plants were verified through Western blot analysis. Dcp1oe was present in the background of Col-0. (e) The protein extracts of CPoe/Dcp1oe and Dcp1oe Arabidopsis leaves were extracted with IP lysis buffer containing 100 μM MG132 and 10 mM DTT and precipitated with Protein A/G Flag antibody beads. CPoe, a transgenic plant expressing CP with a GFP tag. Similar amounts of Dcp1 precipitated by the antibody beads were used for analysis by immunoblotting using anti-Ub and anti-FLAG antibodies.

Given that CP-mediated degradation of Dcp1-Flag can be blocked by the 26S proteasome inhibitor, the subsequent objective was to ascertain whether CP affects AtDcp1 ubiquitination. To this end, we immunoprecipitated Dcp1-Flag with the Flag antibody and subsequently quantified the amount of Dcp1 and ubiquitinated Dcp1 in the immunoprecipitation products using Flag and Ub antibodies, respectively. Figure 5b illustrates a notable elevation in the poly-ubiquitination of Dcp1-Flag by CP in conjunction with an extended co-expression period, accompanied by a decline in the abundance of Dcp1-Flag protein. The poly-ubiquitination of Dcp1-Flag by CP was found to be significantly enhanced when co-expressed with Ub-HA and treated with MG132 for an additional 12 hours following 60 hours of incubation (Fig. 5c).

To determine whether CP affects AtDcp1 ubiquitination in Arabidopsis, we generated Dcp1 overexpression transgenic plants (Dcp1oe) and CP-GFP/Dcp1oe plants by crossing transgenic Dcp1oe plants with transgenic CP-GFP plants (Liu et al., 2023). The presence of Dcp1 in Dcp1oe or both transgenes in the F2 generation of CP-GFP/Dcp1oe heterozygous plants was confirmed through genotyping and Western blot analysis. As illustrated in Fig. 5d, the levels of AtDcp1 protein were significantly lower in CP-GFP/Dcp1oe transgenic plants compared to Dcp1oe, which is consistent with the results obtained from *N. benthamiana*. The phenotype of Dcp1oe was found to be comparable to that of Col-0, while and CP-GFP/Dcp1oe exhibited obvious developmental defects in leaves (Fig. S4), displaying downwardcurled leaf margins similar to the CP-GFP over-expression transgenic plants (Liu et al., 2022). Poly-ubiquitination was markedly elevated in CP-GFP/Dcp1oe transgenic Arabidopsis relative to Dcp1oe plants (Fig. 5e). This result corroborates the hypothesis that CP mediates AtDcp1 degradation via the 26S proteasome pathway *in vivo*.

Given the AtDcp1 protein exhibited elevated accumulation in plants treated with 3-MA in comparison to those treated with DMSO (Fig. S3a). A further experiment was conducted to ascertain whether the reduction of AtDcp1 mediated by CP can also be prevented by the autophagy inhibitor 3-MA. As illustrated in Figure 6a, the autophagy inhibitor 3-MA (10 mM) was observed to block CP-mediated degradation of Dcp1-Flag. These results indicate that CP-mediated degradation of Dcp1 is likely also degraded by autophagy in plant cells. To substantiate this hypothesis, we employed mCherry-tagged N. benthamiana ATG8a (RFP-ATG8a) as an autophagosome marker to monitor autophagy (Li et al., 2017). The observation of numerous punctate RFP fluorescent structures, presumed to represent pre-autophagosome or autophagosome structures (Fig. 6b, top panel and 6c), was made when Dcp1 was transiently overexpressed with CP-GFP in *N. benthamiana* plants. However, the structure is barely discernible when Dcp1 is transiently overexpressed in the presence of pG1300 with free GFP (Fig. 6b, bottom panel), and the structure was not observed when Dcp1 was transiently overexpressed with CP-GFP or pG1300 in the absence of RFP-ATG8a (Fig. S5) as well. These results lend support to the hypothesis that CP mediates AtDcp1 degradation via the autophagy pathway in *vivo*.

**FIGURE 6.**
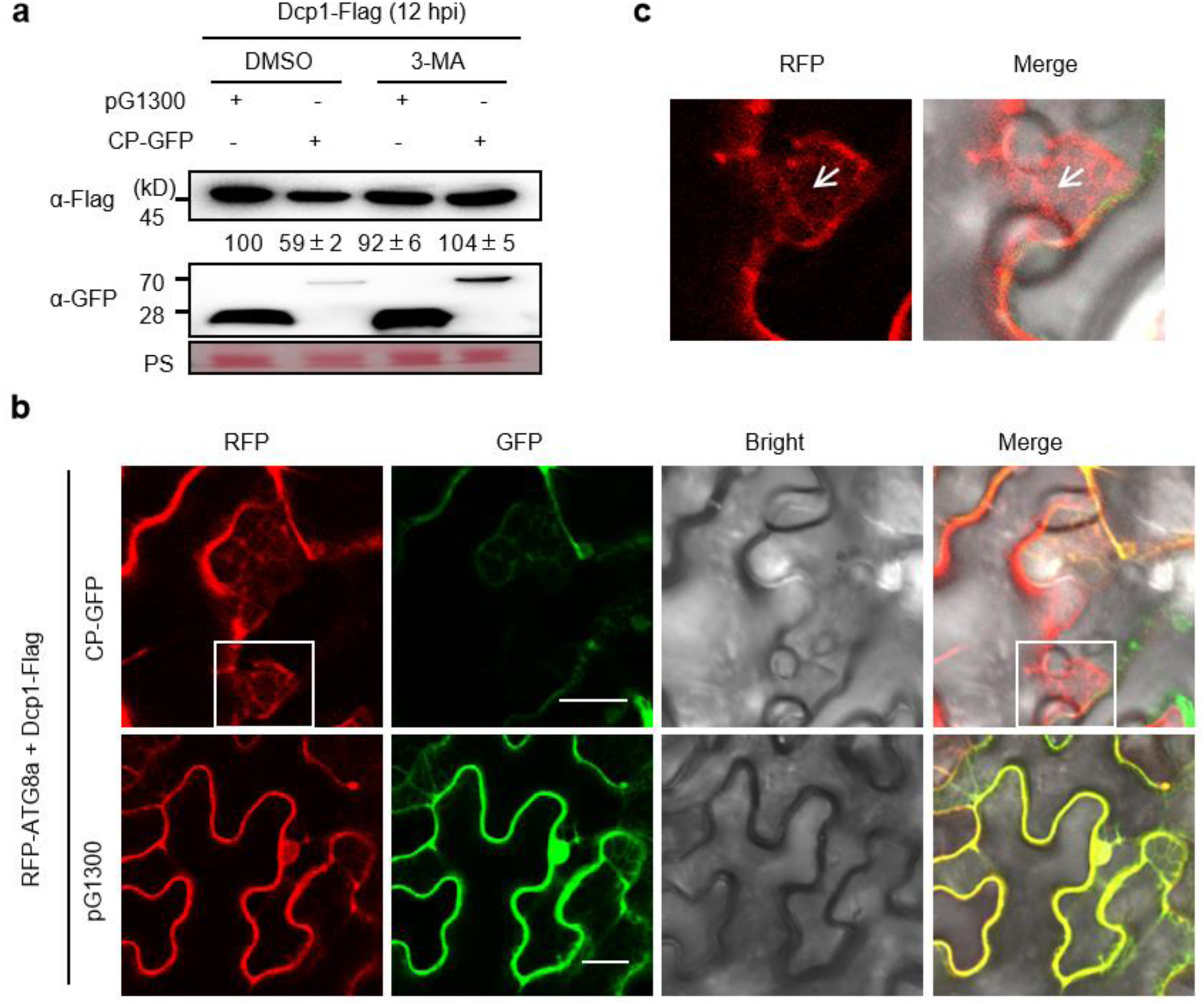
CP mediates AtDcp1 degradation via autophagy pathway. (a) CP-triggered AtDcp1 degradation was blocked by autophagy inhibitor 3-MA. Dcp1-Flag was co-expressed with or without CP-GFP in *N. benthamiana* by agroinfiltration and treated with 3-MA (10 μM) or DMSO at 60 hpa. The extracts were isolated after treatment with 3-MA for western blotting at 12 hpa. PS was used as a loading control to monitor input protein amounts. (b) Micrographs showing cells expressing mCherry-ATG8a with Dcp1-Flag in presence of GFP (pG1300) or CP-GFP on *N. benthamiana plants*. Infiltrated *N. benthamiana* leaves were examined at 48 hpa. Obvious pre-autophagosome or autophagosome structures (boxed) were observed in infiltrated leaves expressing mCherry-ATG8a with Dcp1-Flag in presence of CP-GFP at 2 dpi. Bars represent 20 μm. (c) Enlarged image of the figure (b) in the box. Arrows indicate autophagic structures.

### CP suppresses AtDcp1 function to stabilize the known NMD targets and facilitate viral infection

To investigate whether CP-mediated degradation of AtDcp1 impedes its function, three transcripts, *At1G01060* (Garcia et al., 2014), *At5G64430,* and *At5G22570* (Rayson et al., 2012), targeted by nonsense-mediated decay (NMD), were analyzed in Col-0, Dcp1oe homozygous, F2 CP-GFP/Dcp1oe heterozygous, and *dcp1-4* mutant plants. The results demonstrate a significant increase in mRNA accumulation in *At1G01060*, *At5G64430*, and *At5G22570* in Arabidopsis *dcp1-4* mutant plants, and a decrease in Dcp1oe transgenic plants, as illustrated in Fig. 6a. The mRNA levels of *At1G01060*, *At5G64430*, and *At5G22570* were found to be reduced in Dcp1oe transgenic plants. However, co-overexpression of CP restored these levels, indicating that CP-mediated degradation of AtDcp1 hinders its function, thereby stabilizing previously established NMD targets.

To ascertain whether CP-mediated degradation of AtDcp1 facilitates viral infection, we conducted a mechanical inoculation of four-week-old seedlings of the Col-0, Dcp1oe homozygous, and F2 CP-GFP/Dcp1oe heterozygous plants with in vitro transcribed infectious RNA from CPB and maintained them at 18℃. As illustrated in Fig. 6b, the northern blot results indicate that the CPB RNAs were significantly lower in the Dcp1oe plants than in their wild-type counterparts. The reduction in CPB RNA levels resulting from AtDcp1 overexpression was reversed in heterozygous F2 CP-GFP/Dcp1oe plants. These findings indicate that CP combats RNA decay by targeting AtDcp1 for degradation via the host 26S proteasome and autophagy pathways, thereby facilitating viral infection.

## DISCUSSION

Accumulated studies have shown that RNA decay plays a crucial role in plant defense responses against viral infections by impacting viral mRNA stability (Li and Wang, 2019; Mailliot et al., 2022; Wu et al., 2023a). PBs, which are RNA decay sites, have been associated with infection by several RNA viruses (Pérez-Vilaró et al., 2012; Ariumi et al., 2011; Beckham et al., 2007). In this study, we observed a significant increase in the number and size of PB markers following TCV infection. The markers were translational fusions to GFP fluorescent reporters in 14-day post-infection *N. benthamiana* systemic leaves (Figs 1b & 1c); however, free GFP remained unaffected. Our findings are in agreement with previous reports on TMV and CaMV infections, which also demonstrated a marked increase in the size and number of PB foci in systemic leaves (Conti et al., 2017; Hoffmann et al., 2022). Moreover, the level of viral RNA in Arabidopsis *dcp1-4* mutant plants infected by CPB, which were TCV mutants with compromised VSR, was markedly higher than that observed in wild-type plants (Fig. 1d). Conversely, TCV infection was suppressed by ectopic overexpression of AtDcp1 from Arabidopsis (as demonstrated in Fig. 7b). These findings are consistent with previous reports indicating that the CaMV viral transactivator protein (TAV) binds and destabilizes the RNA decapping complex VCS, resulting in the buildup of NMD substrates and the virus within the cell (Lukhovitskaya and Ryabova, 2019). These results agree with prior reports in which XRN4 was identified as a key factor in limiting virus accumulation within *N. benthamiana* or Arabidopsis plants via the RNA decay pathway during infections of rice stripe virus, TBSV, CNV, TuMV, TMV, and TCV (Cheng et al., 2007; Jaag and Nagy 2009; Peng et al., 2011; Jiang et al., 2018; Wu et al., 2023b). These findings confirm that RNA decay serves as an antiviral mechanism in Arabidopsis plants.

**FIGURE7.**
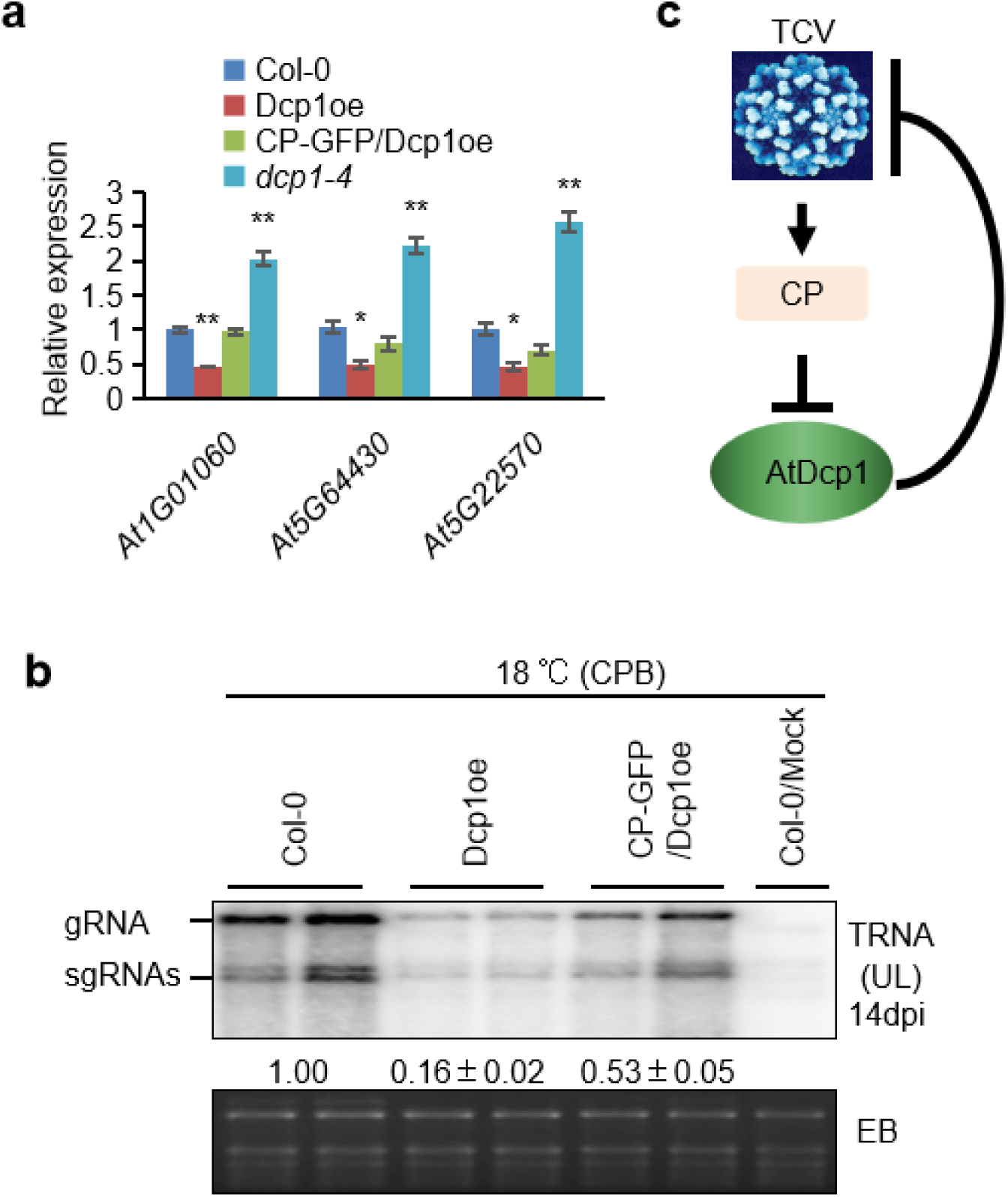
TCV CP blocked AtDcp1 function to facilitate virus replication. (a) Quantitative RT-PCR analysis of known NMD target transcripts containing premature termination codons (PTCs; *At1G01060* (Garcia *et al*., 2014) and unknown NMD triggering signal (*AtG22570*) (Rayson *et al*., 2012) in wild type, transgenic line Dcp1oe, F2 generation plants of CP-GFP/Dcp1oe and *dcp1* mutant Arabidopsis plants. Expression was normalized against AtActin1 transcripts, which served as an internal standard. Each mean value was derived from three independent experiments (n = 3 samples). Values represent the mean ± standard deviation (SD). Asterisks indicate a significant difference (*P < 0.05; **P < 0,01; Student’s t test). (b) Northern blot hybridization of the total RNAs extracted from the UL of CPB-infected Arabidopsis Col-0, transgenic plants Dcp1oe and CP-GFP/Dcp1oe plants. Each RNA sample was extracted from six ULs pooled from six plants. The probe was DIG-labeled DNA, which was a 667-nt fragment of TCV. EB: ethidium bromide-stained Northern gel. See Wu *et al*. (2021) for a detailed procedure. (c) Model of the interaction between the capsid protein of TCV and Dcp1 of the mRNA decay pathway.

Viruses are intracellular parasites that require host cells for reproduction. Plant and animal viruses use a range of strategies in the molecular arms race with host cells to evade RNA decay (Popp et al., 2020). Potyviral HC-Pro, genome-linked protein (VPg), CaMV TAV, and CP can function as suppressors of RNA decay by targeting 5′ to 3′ RNA decay components, including XRN4, Dcp2, or VCS (Li and Wang, 2018; Lukhovitskaya and Ryabova, 2019; Wu et al., 2023b). Pea enation mosaic virus 2 (PEMV2) p26 protein antagonizes NMD in the cytoplasm of infected cells by stabilizing NMD target transcripts, particularly those with lengthy, structured, and GC-rich 3′ UTRs (May *et al*. 2020). The BYSMV P protein ensures the assembly of genuine viral RNA-protein complexes by recruiting a conserved mRNA decay enzyme, host carbon catabolite repression 4, resulting in the degradation of N-bound cellular mRNAs (Zhang et al., 2020). The Dcp1/Dcp2 complex is essential for RNA decay, it has been reported that TuMV VPg disrupts the interaction between NbDcp1 and NbDcp2 by relocating NbDcp2 from the cytoplasm to the nucleus, which can impede viral RNA degradation through RNA decay (Li and Wang, 2018). Our research shows that co-infiltrated YN-Dcp2 and Dcp1-YC with CP reconstituted YFP signals at 2 dpa (Fig. 3). This indicates that, unlike TuMV VPg, CP does not hinder the formation of cytoplasmic Dcp1/Dcp2 granules, even though it interacts with AtDcp1 (Fig. 2). This finding proposes that various viruses may have developed distinctive mechanisms to avoid RNA decay during their long-term evolutionary process (Guo *et al*., 2018; Li and Wang, 2019; Popp *et al*., 2020; May et al., 2021; Wu et al., 2023a).

Over the past decade, numerous studies have highlighted the intricate interplay of UPS and autophagy in the relationship between plants and viruses. Despite operating as a defense mechanism against viruses, both UPS and autophagy can be exploited by viruses to promote their infection in plants (Wu et al., 2023a; Sun et al., 2023). In addition to directly hijacking or suppressing proteins associated with UPS through viral proteins (Jia et al., 2016; Li et al., 2019; Liu et al., 2021; Zhang et al., 2011; Yang et al., 2018; Cao et al., 2023; Yang et al., 2022b; Yang et al., 2022c; Yang et al., 2023; Niu et al., 2022; Sun et al., 2022), viruses can modulate RNA levels of genes related to UPS and autophagy to counteract antiviral defense (Mei et al., 2020; Lai et al., 2009; Chen et al., 2020; Fu et al., 2022; Wang et al., 2023). Additionally, some plant RNA viruses have evolved to encode proteins that exploit cellular UPS and autophagy to degrade specific host defense components and facilitate viral replication. For instance, the degradation of SGS3 proteins, which compromise of RNA silencing-mediated antiviral immunity via UPS and autophagy (Cheng et al., 2016; Tong et al., 2021). A plant-specific, membrane-associated protein known as REM, in its S-acylated form, displays antiviral defense properties. RSV infection hinders the S-acylation of REM, which then leads to the degradation of non-S-acylated REM through autophagy (Fu et al., 2018). Wang et al. (2022) reported that the interaction between plant UVRAG and ATG14 plays a pivotal role in regulating autophagosome maturation and other aspects of geminiviral infection. The study indicates that the presence of CP leads to a decrease in the abundance of AtDcp1 (Fig. 4), while an increase in the poly-ubiquitination of Dcp1-Flag (Figs 5b, 5c & 5e) and pre-autophagosome or autophagosome structures (Fig. 6b top panel and 6c) suggesting CP’s role in mediating AtDcp1 degradation via both 26S proteasome and autophagy pathways. These findings reveal a molecular mechanism underlying how the CP deploys host 26S proteasome and autophagy pathways to downregulate the expression level of AtDcp1.

In this study, a reduction in mRNA levels of NMD target transcripts and virus accumulation was observed upon the overexpression of AtDcp1. However, this negative effect was offset by the expression of CP (Fig. 7b). Therefore, CP as a highly efficient VSR (Qu *et al*., 2003) and also disrupts RNA decay by degrading AtDcp1. Consequently, it facilitates viral infection by inhibiting RNA decay. This illustrates how a viral protein can selectively target RNA decay signaling components to subdue host cellular antiviral RNA decay defense and uphold viral infection in plants. We propose that TCV, via the capsid protein, has evolved a strategy to lower Dcp1 levels and dampen antiviral activities of RNA decay, which in turn contributes to viral infection (Fig. 7c).

## MATERIALS AND METHODS

### Plant materials and TCV variants

All *A. thaliana* and *N. benthamiana* plants were grown in pots in a growth chamber at 23 ℃ with 14 h of daylight and 60% humidity. The *A. thaliana* line *dcp1-4* (SALK_014408C) mutant was provided by Dr. Feng Qu at Ohio State University. Homozygous mutation lines were screened by using primers as listed in Supplementary Table S2. The CP-GFP and CPC-GFP Arabidopsis transgenic plants used in this study have been described previously (Liu et al., 2022). The Arabidopsis plants were inoculated with *in vitro*-transcribed viral RNA at the age of 3 weeks to 4 weeks. After inoculation, the infected plants were moved into a versatile environmental test chamber (SANYO) at 18 ℃ under a 16-h/8-h photoperiod, 60% humidity, and light intensity of 160–190 μmol/m^2^/sec. Arabidopsis (ecotype Col-0) plants were transformed via floral-dip method. Transformants were screened in Murashige and Skoog (MS) medium supplied with 20 μg/ml hygromycin B and then further confirmed by PCR.

The infectious clone CPB contained one amino acid change at amino acid residues 130 (R130T) of CP, which was constructed in Dr. Qu’s laboratory at Ohio State University as previously described (Cao et al., 2010).

### TCV infection procedure

For inoculation, *A. tumefaciens* strain GV3101, which contained proper constructs, infiltrated fully expanded leaves from 3-week-old *N. benthamiana* plants at OD600 of 0.3. At 14 dpa, upper uninfiltrated fully expanded leaves were used for infiltration by *A. tumefaciens* strain GV3101, which contained proper constructs.

### Infection of Arabidopsis with *in vitro* transcripts

The *in vitro* transcripts of TCV variants were produced using the Transcript Aid T7 High Yield Transcription Kit (Fermentas, Glen Burnie, MD) in accordance with the manufacturer’s instruction. The integrity and the concentration of the transcripts were examined using agarose gel electrophoresis. The inoculum was prepared by diluting the transcripts to 10 ng/μL by using the inoculation buffer (pH 9.2) containing 50 mM glycine, 30 mM K_2_HPO_4_, 1% bentonite, and 1% celite. 10 μL inoculum was used for mechanical inoculation with a gloved finger on each *Arabidopsis* leaf. Usually for each mutant and the wild type controls, at least six plants were infected with the infectious transcripts of clones described above, on three fully expanded leaves per plant.

### Northern blot assay

For RNA blot analysis, total RNAs were isolated from the virus-inoculated Arabidopsis leaves (IL) at 5 dpi or from the upper uninoculated Arabidopsis leaves (UL), which were approximately 1 cm long at 14 dpi by using the TRIzol reagent (Tiangen Biotech Beijing Co., Ltd.), following the manufacturer’s instruction. To minimize sampling errors, we pooled leaves from six different plants (one leaf per plant) before RNA extraction. Different amounts of total RNA from Arabidopsis were used for Northern blot assay to detect vsRNAs (5 μg) and viral RNA (1 μg) according to published protocols (Zhang et al., 2012; Wu et al., 2021).

### Plasmid construction

For the Y2H assay, the full-length coding sequence of Arabidopsis *Dcp1* (*AT1G08370.1*) was amplified from *A. thaliana* cDNA by RT-PCR using the primers (Supplementary Table S3) introducing appropriate restriction sites, followed by insertion of the resulting fragment into the pGBKT7 vector. AD-CP, CPB and CPC which contained the full-length coding sequences of CP, CPB, and CPC were constructed as previously described (Liu et al., 2022).

For the BiFC and subcellular experiments, the full-length *AtDcp1* was amplified from the obtained yeast pGBKT7 construct mentioned above with appropriate nucleotide primers (Supplementary Table S3) and then was subcloned in-frame with the binary vector p2YC or pG1300 to obtain constructs Dcp1-YC and Dcp1-GFP respectively. The full-length coding sequence of and *Dcp2* (*AT5G13570.2*) was amplified from *A. thaliana* cDNA by RT-PCR using the primers (Supplementary Table S3) introducing appropriate restriction sites, followed by insertion of the resulting fragment into pYN1 or pG1300 vector, resulting in YN-Dcp2 and Dcp2-GFP, respectively. p2YC, pYN1, YN-CP, CPB and CPC were constructed as previously described (Liu et al., 2022). Plasmids for subcellular localization, including CP-RFP, CPB-RFP, and CPC-RFP, were constructed as previously described (Liu et al., 2022).

For protein in *vitro* degradation assays, the full-length sequences of TCV CP and AtDcp1 were amplified by PCR using specific primers, and the constructed into the pet28(a)-GFP and pet28(a)-sumo-Flag vector using BamHI and SalI restriction sites, resulting in pet28(a)-GFP-CP with GFP-HIS tagged fusion proteins and pet28(a)-sumo-Flag-Dcp1 with Flag-tagged fusion protein respectively.

To generate the AtDcp1 overexpressing transgenic lines, we constructed the binary expression vector Dcp1-Flag with Flag tag by using appropriate nucleotide primers (Supplementary Table S3) by accommodating the cDNAs of AtDcp1 in pF1300. pF1300 was constructed by replacing the GFP of pG1300 with a synthetic Flag epitope tag, designated as Flag *Bam*HI and *Sac*I (Supplementary Table S3). Both constructs were under the control of CaMV35S promotor.

Additional details of all vectors and amplification primers are available upon request. The construct nucleotide sequences were validated via sequencing.

### Y2H assay

A Gal4-based Y2H system was used to detect the interaction between various viral proteins and Arabidopsis proteins. Approximately 100 μL of freshly prepared *Saccharomyces cerevisiae* (strain AH109) competent cells was cotransformed with the BD and AD plasmids with the help of 5 μL of herring carrier DNA. Transformants were uniformly plated on agar-solidified double drop out (DDO) medium SD/−Leu/−Trp (SD-LW). Plates were incubated in a constant-temperature incubator at 30 ℃. Transformants were identified based on PCR amplification. Protein–protein interactions were detected by transferring yeast cotransformants that were grown on the DDO medium to plates containing the agar-solidified QDO medium SD/−Leu/−Trp/−His/−Ade (SD-LWHA).

### BiFC, subcellular localization and co-localization analysis

For the BiFC assays, complementary vectors containing target genes for interaction identification were co-agroinfiltrated into *N. benthamiana* leaves. At 2 d post-infiltration, the yellow fluorescence, peroxisome signals (CFP), and 4′,6-diamidino-2-phenylindole (DAPI)-stained cell nuclei were detected under confocal laser-scanning microscope. For the subcellular localization and co-localization analysis the fluorescence of the corresponding plasmid was visualized in transient *N. benthamiana* leaves as previously described (Lin et al., 2022).

### Transient protein expression and confocal microscopy

*A. tumefaciens* strain GV3101, which contain proper constructs, infiltrated fully expanded leaves from 3-week-old *N. benthamiana* plants. For co-agroinfiltration and BiFC experiments, the A. tumefaciens suspensions were mixed in 1:1 ratio. For western blot and co-immunoprecipitation (Co-IP) assays, *A. tumefaciens* cells harbouring fusion protein were cultured to an OD600 of 0.6–0.8 to optimize protein expression. For confocal microscopy analysis, the plant samples expressing recombinant proteins were imaged using an Olympus FV1000 confocal laser scanning microscope available through the Institute of Tropical Bioscience and Biotechnology, Chinese Academy of Tropical Agricultural Sciences and according to manufacturer’s protocol. DAPI, YFP, and RFP were excited using 405-, 514-, and 558-nm laser lines, respectively. Quantification of fluorescence intensity was performed using IMAGEJ (http://rsbweb.nih.gov/ij/).

### DAPI staining of plant nuclei

The infiltrated leaf patches were immersed in a DAPI solution (4′,6-diamidino-2-phenylindole dihydrochloride; 15 μg/ml in agroinfiltration buffer) and incubated in the dark for 1 to 2 hours before examination of leaf sections using epifluorescence microscopy.

### Protein extraction and Western blot analysis

Protein was extracted from infiltrated leaf patches of *N. benthamiana* leaf pool by using RIPA buffer (10 mM Tris-HCl at pH 8.0 with 1 mM EDTA, 0.5 mM EGTA, 1% Triton X-100, 0.1% sodium deoxycholate, 0.1% SDS, 140 mM NaCl, 1 mM PMSF, and 1X ProBlockTM Gold Plant Protease Inhibitor Cocktail; the last two reagents should be added immediately before use). Western blot analysis was carried out as previously described (Wu et al., 2023b). Anti-Flag, Anti-HA and anti-GFP antibodies were purchased from Proteintech (Wuhan, China). Anti-Ubiquitin (affinity purified) was purchased from Agrisera, Blotted membranes were washed thoroughly and visualized using ImageQuant LAS 4000mini, according to the manufacturer’s protocol (ECL; GE Healthcare).

### Co-Immunoprecipitation (Co-IP) assays

For Co-IP assays, the different combinations were co-expressed in *N. benthamiana* leaves. After infiltration for 40-48 h, the tobacco leaf tissues were harvested and ground into powder with liquid nitrogen and used for Co-IP assays. The native total protein was extracted using IP lysis buffer (Thermo Scientific) with 10 mM DTT, 1× EDTA-free protease inhibitor cocktail (Roche, Basel, Switzerland). After incubation for 30 min at 4 °C, the homogenate was centrifuged at 12000 g, 4 °C for 10 min, and the process repeated. The supernatant was incubated with 20 μL Protein A/G Flag antibody beads for approximately 2 h at 4 °C with gentle shaking. Protein beads were prewashed three times with 1×PBS. After co-incubation, the immunoprecipitates were washed three times with 1×PBS and resuspended in 50-100 μL 2×SDS-PAGE sample buffer containing 500 mM Tris-HCl, 50% glycerin, 10% SDS, 1% bromophenol blue and 2% β-mercaptoethanol, pH=6.8). Subsequently, the proteins were analyzed by anti-FLAG/ GFP (Proteintech) / Ubiquitin (Agrisera) antibody.

### Chemical treatments

For the protein degradation assay, infiltrated leaves were treated with PBS buffer containing 2% dimethyl sulfoxide (DMSO; control) or an equal volume of DMSO with 5 mM 3-MA (Sigma) for inhibition of autophagy, or an equal volume of DMSO with 100 μM MG132 (Sigma-Aldrich)) for inhibition of the 26S proteasome for 12 h at 4 d post infiltration (dpi).

### Recombinant protein expression

The recombinant plasmid pet28(a)-GFP-CP, pet28(a)-sumo-Flag-Dcp1 or empty vector pet28(a)-GFP was transferred into Escherichia coli BL21 (DE3) cells (WEIDI Biotech, Shanghai, China) and cultured in LB broth supplemented with kanamycin (50 μg/ml) at 37°C, shaking at 200 rpm. The overnight cultures were diluted 1/10 with fresh LB broth with appropriate antibiotics, the culture was incubated at 37°C until the OD_600nm_ reached 0.5 - 0.6, followed by induction using 0.1-1 mM IPTG for an additional 4 h (37 °C, 200 rpm). Cells were harvested via centrifugation (20,000× g, 10 min, 4 °C) and then the pellet was resuspended in lysis buffer (50 mM Tris, 5% glycerol (v/v), and 50 mM NaCl; pH 7.5), and sonicated for 15 min with a cycle of 10 s pulse and 20 s pause. The sonicated sample was centrifuged and the supernatant was kept for in vitro protein degradation assay. Furthermore, western blotting was performed to confirm the band of recombinant protein observed on SDS-PAGE.

### *In vivo* and *in vitro* protein degradation assay

For protein in vitro degradation assays, the cell extract of Flag-Dcp1 was mixed with that of GFP-CP or GFP extract at a ratio of 1:1, and incubated at 25°C. The incubated protein extracts were sampled at four time points, and the reactions were stopped by addition of SDS sample buffer and boiled for 5 min. Then, western blot analyses were conducted as described above.

For protein in *vivo* degradation assays, Dcp1-Flag was transiently co-expressed with PBS buffer or GFP or CP-GFP in *N. benthamiana* leaf cells, respectively. Dcp1-Flag and GFP or CP-GFP were co-agroinfiltrated into *N. benthamiana* leaves with or without 3-MA or MG132 treatments and analyzed by western blotting.

### RNA extraction, RT-qPCR analysis

Total RNA was isolated using TRIzol and treated with DNase I (mona) following the manufacturer’s instruction. cDNA synthesized from the reverse transcription of RNA samples was used to quantify TCV accumulation levels and determine the mRNA levels of target genes. The cDNA synthesis, RT-qPCR assays and ddPCR analysis were conducted as described previously (Zhang et al., 2012; Liu et al., 2022). The expression of the AtActin1 gene was referred as an internal control to normalize cDNA concentrations. Information about all the primers used is summarized in Supplementary Table S1. All RT-qPCRs were carried out in three independent biological replicates and triplicate for each cDNA sample.

### Statistical analysis

Unless otherwise stated, all experiments were performed with at least three biological replicates in all cases. Significant differences between samples in gene expression were statistically analyzed with Student’s tests in GraphPad Prism 8.0 software. Values of p < 0.05 and p < 0.01 were taken as statistically significant.

## DATA AVAILABILITY

The data that support the findings of this study and the materials used during the study are available from the corresponding author on reasonable request.

## AUTHOUR CONTRIBUTIONS

XC-Z: conceptualization; KX-W, QX-X, XT-L, Y-F, SX-L, XL-Y, WB-L and PJ-Z investigation; All authors interpreted data; KX-W: writing - original draft; XC-Z: writing - review & editing; YL-R, MB-R and XC-Z: supervision; XC-Z, and Y-F: funding acquisition; All authors read and approved the final manuscript.

## ACKNOWLEDGEMENTS

We are grateful to Professor Feng Qu (Ohio State University, USA) for providing us with mutant plant lines and plasmids. We are very grateful for the comments and suggestions provided by the anonymous reviewers who reviewed the first version of the manuscript. We thank the Public Technology Research and Sharing Center of the Institute of Tropical Bioscience and Biotechnology Chinese Academy of Tropical Agricultural Sciences for sharing their equipment and providing technical support.

This work was supported by the National Natural Science Foundation of China (32070172) and Natural Science Foundation of Hainan Province (322CXTD528, 324QN352).

---

**Figure S1.**
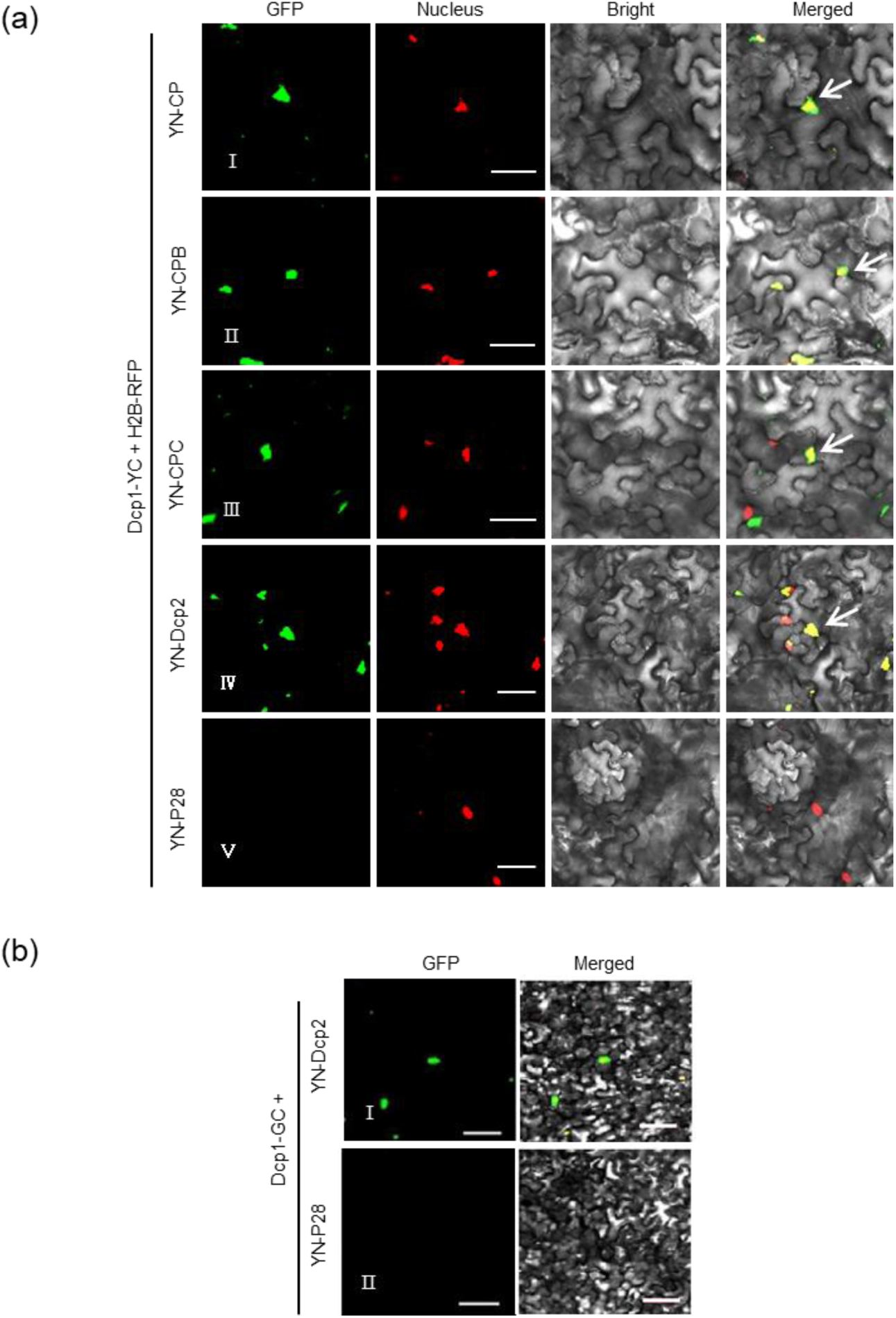
Bimolecular fluorescence complementation (BiFC) assay for protein–protein interactions (a) between TCV CP (I) or its mutant CPB (II) or CPC (III) and AtDcp1. TCV CP, CPB, and CPC were fused with the N-terminal, while AtDcp1 was fused with the C-terminal half of YFP, designated as YN-CP, YN-CPB, YN-CPC and Dcp1-YC, respectively, and transiently expressed in *N. benthamiana* leaves. H2B-RFP was co-infiltrated as a nuclear marker. Fluorescence was monitored by confocal microscopy at 2 dpa. Bar, 50 μm. White arrows represent interaction body colocalized with nucleus. Positive and negative control for bimolecular fluorescence complementation (BiFC) assay.

**Figure S2.**
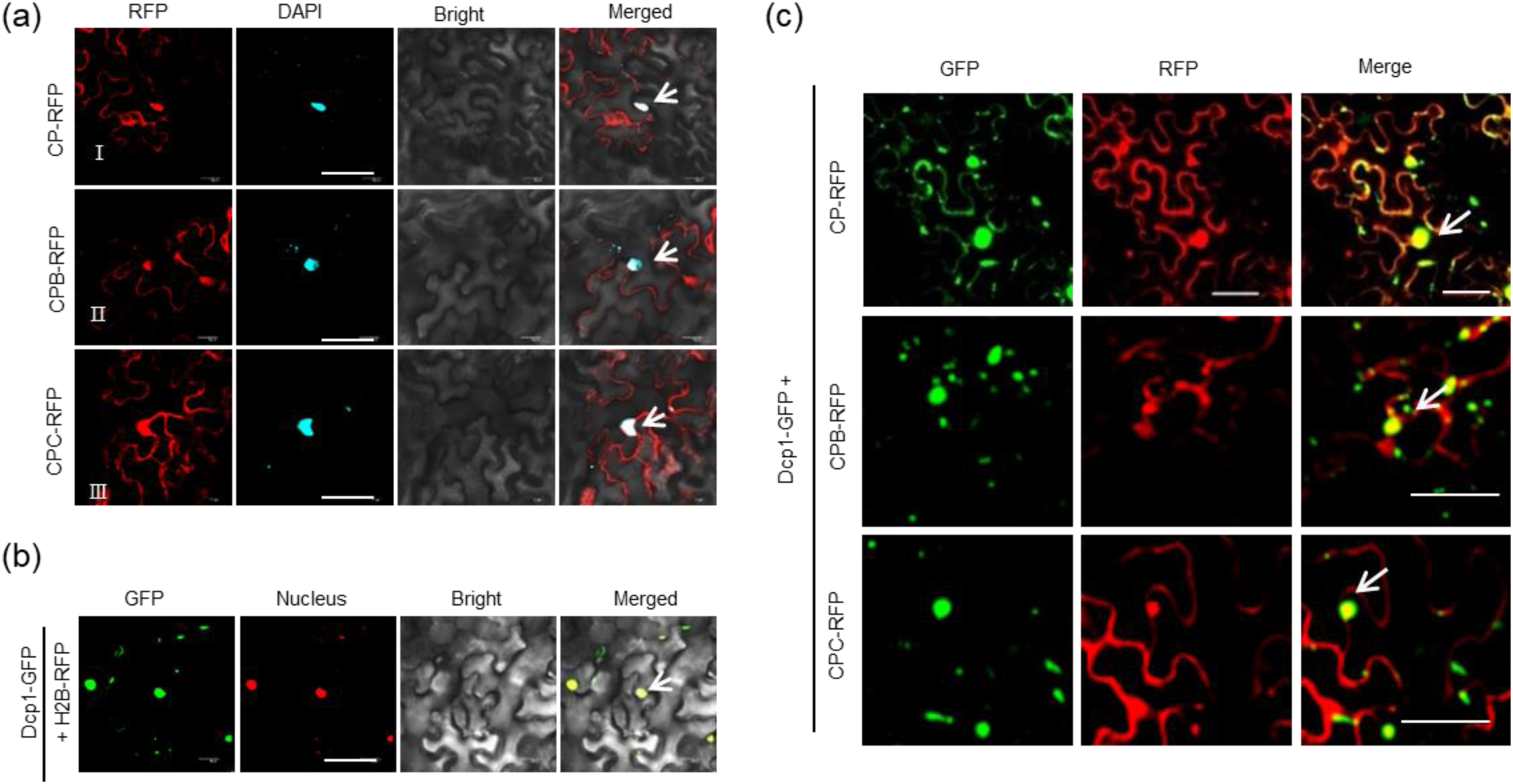
AtDcp1 colocalized with CP, CPB and CPC. (a) Subcellular localization of mCherry fused to CP (CP-RFP), CPB (CPB-RFP), or CPC (CPC-RFP). Bar, 50 μm. White arrows represent the nucleus. (b) Subcellular localization of GFP fused to AtDcp1 (Dcp1-GFP). Bar, 50 μm. White arrow represents the nucleus. (c) Confocal micrographs of *N. benthamiana* co-expressing (CP-RFP) (Ⅰ), CPB (CPB-RFP) (Ⅱ) or CPC (CPC-RFP) (Ⅲ) with Dcp1-GFP at 48 hpa. Bar, 50 μm. White arrows represent nuclear interaction body, respectively.

**Figure S3.**
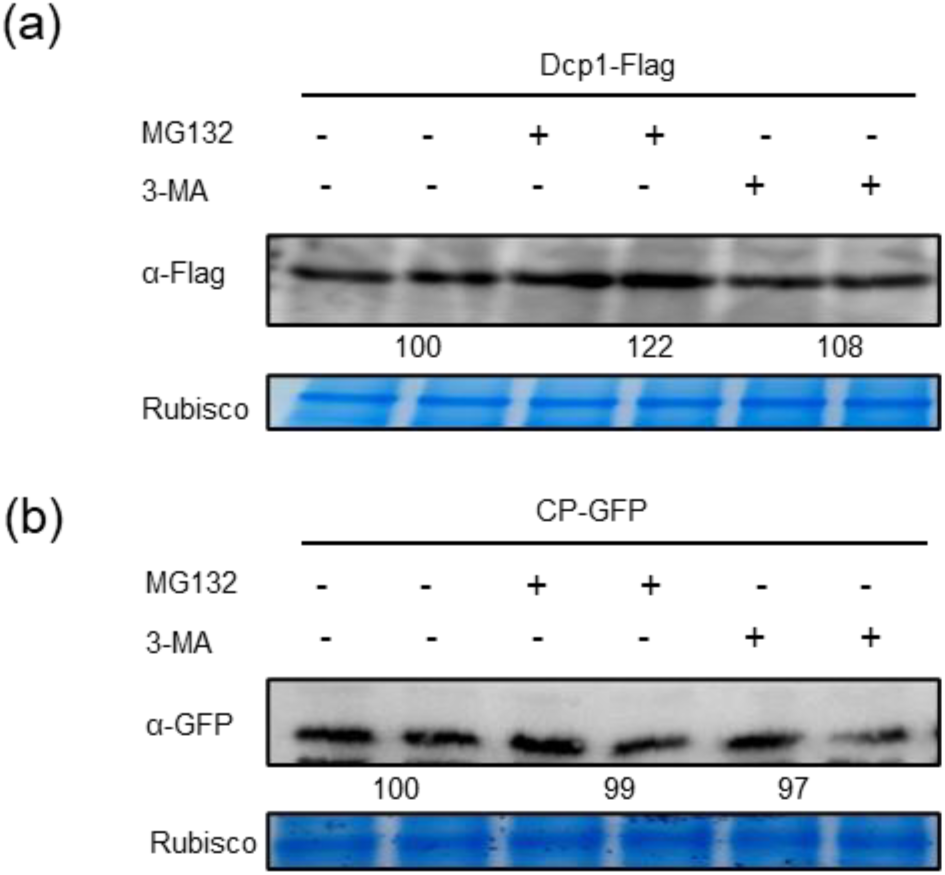
Western blot analysis of Dcp1-Flag (a) or CP-GFP (b) transiently expressed in *N. benthamiana* leaves by *Agrobacterium* infiltration. At 4 dpa, the infiltrated leaf areas were treated with MG132, or 3-MA for an additional 12 h. Total protein was extracted, separated by SDS-PAGE, and detected with Flag antibodies or GFP antibodies. The relative amounts of AtDcp1 or CP are indicated. The bottom panel shows equal protein loading by CBB staining of rubisco.

**Figure S4.**
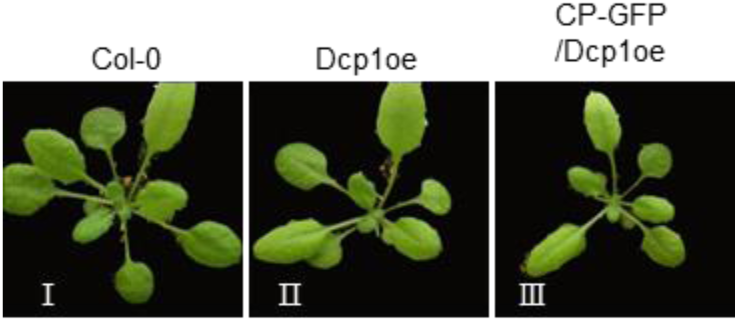
Phenotypes of the Col-0, Dcp1oe, F2 generation CP-GFP/Dcp1oe heterozygous plant. F2 generation CP-GFP/Dcp1oe plants were generated by crossing CP-GFP with Dcp1oe Arabidopsis plants. CP-GFP plants were obtained as described in Liu et al., 2022.

**Figure S5.**
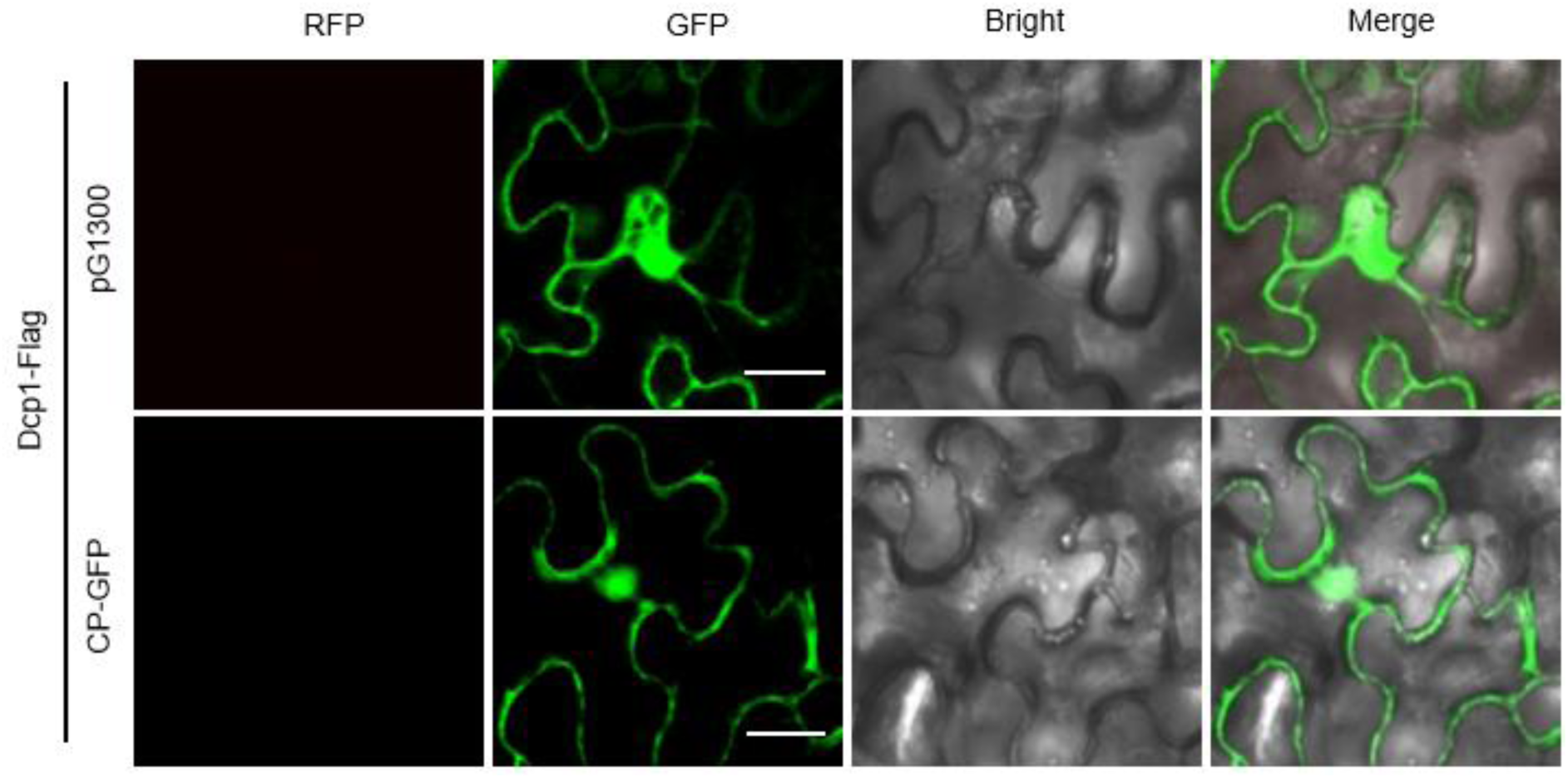
Micrographs showing cells expressing GFP (pG1300) or CP-GFP with Dcp1-Flag in presence of GFP (pG1300) or CP-GFP on *N. benthamiana plants*. Infiltrated *N. benthamiana* leaves were examined at 48 hpa. NO pre-autophagosome or autophagosome structures were observed in infiltrated leaves. Bars represent 20 μm.

**Table S1.**
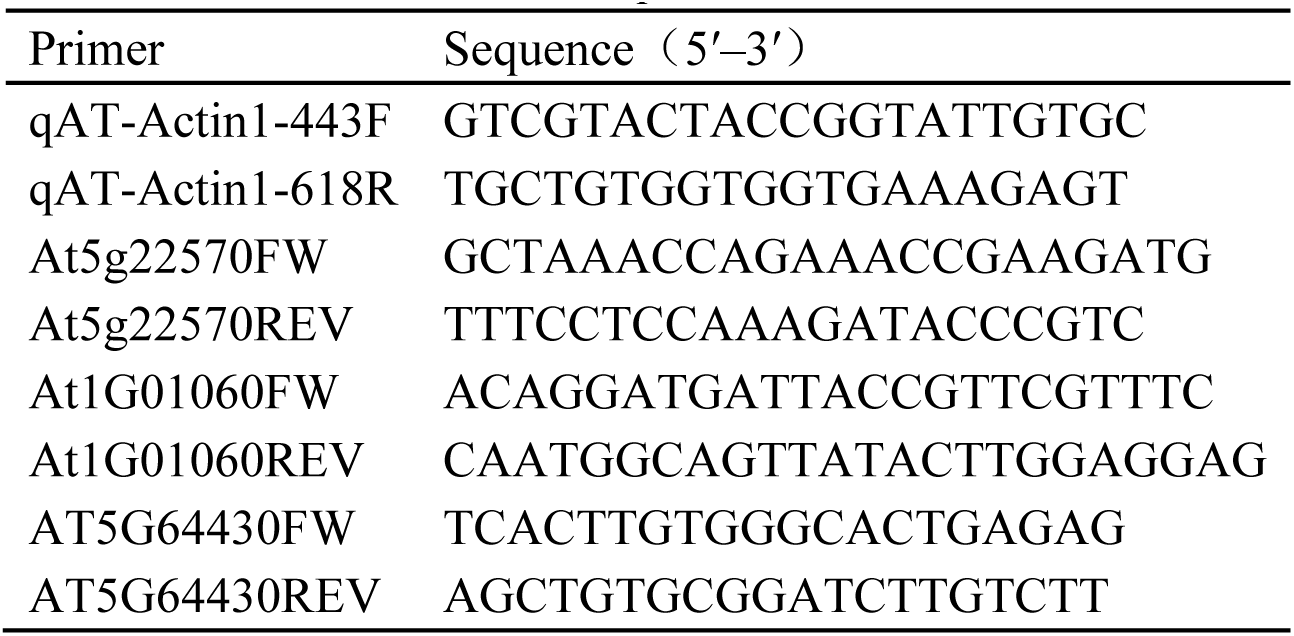
Primers used for the quantitative RT-PCR.

**Table S2.**
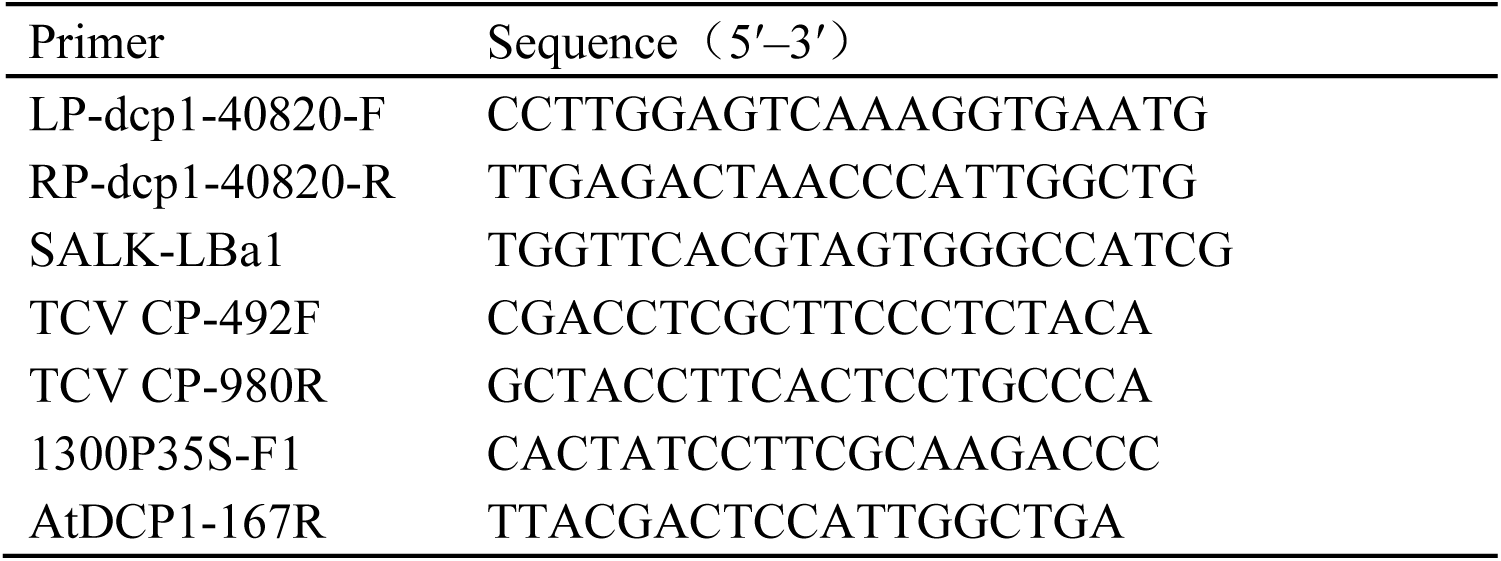
Primers used for Arabidopsis genotyping and transgenic plants.

**Table S3.**
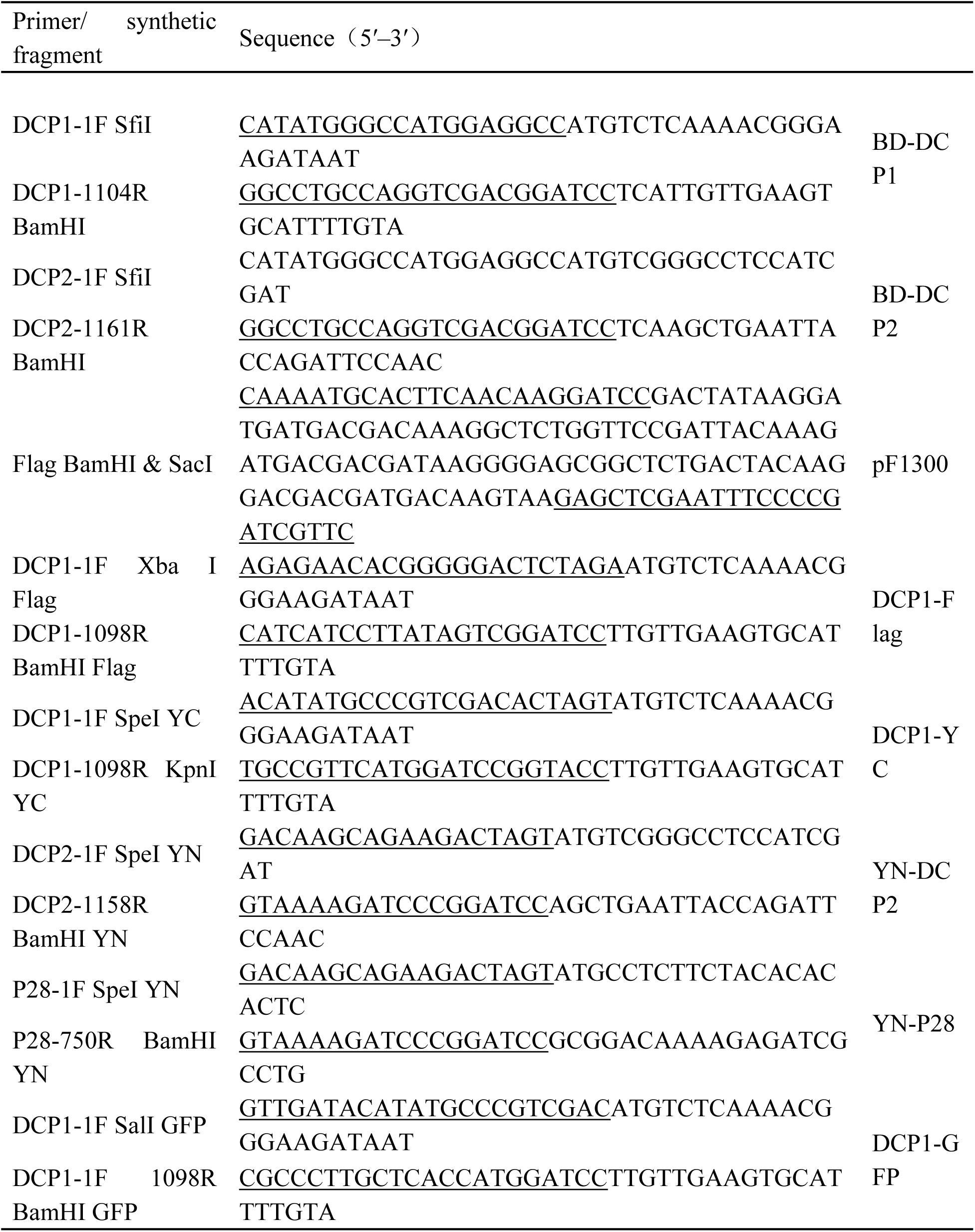
Primers and synthetic fragments used for constructing binary expression. ^a^ compatible adaptor sequences and restriction endonuclease sites are underlined. ^b^ lowercase letters indicate the Flag epitope tag.

## References

Ariumi, Y., Kuroki, M., Kushima, Y., Osugi, K., Hijikata, M., Maki, M., Ikeda, M., & Kato, N. (2011). Hepatitis C virus hijacks P-body and stress granule components around lipid droplets. Journal of virology, 85(14), 6882–6892.

Balistreri, G., Bognanni, C., & Mühlemann, O. (2017). Virus Escape and Manipulation of Cellular Nonsense-Mediated mRNA Decay. Viruses, 9(1), 24.

Beckham, C. J., Light, H. R., Nissan, T. A., Ahlquist, P., Parker, R., & Noueiry, A. (2007). Interactions between brome mosaic virus RNAs and cytoplasmic processing bodies. Journal of virology, 81(18), 9759–9768.

Cao, B., Ge, L., Zhang, M., Li, F., & Zhou, X. (2023). Geminiviral C2 proteins inhibit active autophagy to facilitate virus infection by impairing the interaction of ATG7 and ATG8. Journal of integrative plant biology, 65(5), 1328–1343.

Cao, M., Ye, X., Willie, K., Lin, J., Zhang, X., Redinbaugh, M. G., Simon, A. E., Morris, T. J., & Qu, F. (2010). The capsid protein of Turnip crinkle virus overcomes two separate defense barriers to facilitate systemic movement of the virus in Arabidopsis. Journal of virology, 84(15), 7793–7802.

Chen, Z. Q., Zhao, J. H., Chen, Q., Zhang, Z. H., Li, J., Guo, Z. X., Xie, Q., Ding, S. W., & Guo, H. S. (2020). DNA Geminivirus Infection Induces an Imprinted E3 Ligase Gene to Epigenetically Activate Viral Gene Transcription. The Plant cell, 32(10), 3256–3272.

Cheng, C. P., Jaag, H. M., Jonczyk, M., Serviene, E., & Nagy, P. D. (2007). Expression of the Arabidopsis Xrn4p 5’-3’ exoribonuclease facilitates degradation of tombusvirus RNA and promotes rapid emergence of viral variants in plants. Virology, 368(2), 238–248.

Cheng, X., & Wang, A. (2016). The Potyvirus Silencing Suppressor Protein VPg Mediates Degradation of SGS3 via Ubiquitination and Autophagy Pathways. Journal of virology, 91(1), e01478–16.

Conti, G., Zavallo, D., Venturuzzi, A. L., Rodriguez, M. C., Crespi, M., & Asurmendi, S. (2017). TMV induces RNA decay pathways to modulate gene silencing and disease symptoms. The Plant journal: for cell and molecular biology, 89(1), 73–84.

Derrien, B., Baumberger, N., Schepetilnikov, M., Viotti, C., De Cillia, J., Ziegler-Graff, V., Isono, E., Schumacher, K., & Genschik, P. (2012). Degradation of the antiviral component ARGONAUTE1 by the autophagy pathway. Proceedings of the National Academy of Sciences of the United States of America, 109(39), 15942–15946.

Fu, S., Wang, K., Ma, T., Liang, Y., Ma, Z., Wu, J., Xu, Y., & Zhou, X. (2022). An evolutionarily conserved C4HC3-type E3 ligase regulates plant broad-spectrum resistance against pathogens. The Plant cell, 34(5), 1822–1843.

Fu, S., Xu, Y., Li, C., Li, Y., Wu, J., & Zhou, X. (2018). Rice Stripe Virus Interferes with S-acylation of Remorin and Induces Its Autophagic Degradation to Facilitate Virus Infection. Molecular plant, 11(2), 269–287.

Furuichi, Y., LaFiandra, A., & Shatkin, A. J. (1977). 5’-Terminal structure and mRNA stability. Nature, 266(5599), 235–239.

Garcia, D., Garcia, S., & Voinnet, O. (2014). Nonsense-mediated decay serves as a general viral restriction mechanism in plants. Cell host & microbe, 16(3), 391–402.

Garneau, N. L., Wilusz, J., & Wilusz, C. J. (2007). The highways and byways of mRNA decay. Nature reviews. Molecular cell biology, 8(2), 113–126.

Ge, L., Cao, B., Qiao, R., Cui, H., Li, S., Shan, H., Gong, P., Zhang, M., Li, H., Wang, A., Zhou, X., & Li, F. (2023). SUMOylation-modified Pelota-Hbs1 RNA surveillance complex restricts the infection of potyvirids in plants. Molecular plant, 16(3), 632–642.

Goldberg A. L. (2012). Development of proteasome inhibitors as research tools and cancer drugs. The Journal of cell biology, 199(4), 583–588.

Guo, L., Vlasova-St Louis, I., & Bohjanen, P. R. (2018). Viral manipulation of host mRNA decay. Future virology, 13(3), 211–223.

Hoffmann, G., Mahboubi, A., Bente, H., Garcia, D., Hanson, J., & Hafr N, A. (2022). Arabidopsis RNA processing body components LSM1 and Dcp5 aid in the evasion of translational repression during Cauliflower mosaic virus infection. The Plant cell, 34(8), 3128–3147.

Hogg J. R. (2016). Viral Evasion and Manipulation of Host RNA Quality Control Pathways. Journal of virology, 90(16), 7010–7018.

Hsu, C. L., & Stevens, A. (1993). Yeast cells lacking 5’-->3’ exoribonuclease 1 contain mRNA species that are poly(A) deficient and partially lack the 5’ cap structure. Molecular and cellular biology, 13(8), 4826–4835.

Jaag, H. M., & Nagy, P. D. (2009). Silencing of Nicotiana benthamiana Xrn4p exoribonuclease promotes tombusvirus RNA accumulation and recombination. Virology, 386(2), 344–352.

Ji, M., Zhao, J., Han, K., Cui, W., Wu, X., Chen, B., Lu, Y., Peng, J., Zheng, H., Rao, S., Wu, G., Chen, J., & Yan, F. (2021). Turnip mosaic virus P1 suppresses JA biosynthesis by degrading cpSRP54 that delivers AOCs onto the thylakoid membrane to facilitate viral infection. PLoS pathogens, 17(12), e1010108.

Jia, Q., Liu, N., Xie, K., Dai, Y., Han, S., Zhao, X., Qian, L., Wang, Y., Zhao, J., Gorovits, R., Xie, D., Hong, Y., & Liu, Y. (2016). CLCuMuB βC1 Subverts Ubiquitination by Interacting with NbSKP1s to Enhance Geminivirus Infection in Nicotiana benthamiana. PLoS pathogens, 12(6), e1005668.

Jiang, S., Jiang, L., Yang, J., Peng, J., Lu, Y., Zheng, H., Lin, L., Chen, J., & Yan, F. (2018). Over-expression of Oryza sativa Xrn4 confers plant resistance to virus infection. Gene, 639, 44–51.

Lai, J., Chen, H., Teng, K., Zhao, Q., Zhang, Z., Li, Y., Liang, L., Xia, R., Wu, Y., Guo, H., & Xie, Q. (2009). RKP, a RING finger E3 ligase induced by BSCTV C4 protein, affects geminivirus infection by regulation of the plant cell cycle. The Plant journal: for cell and molecular biology, 57(5), 905–917.

Li, F., & Wang, A. (2018). RNA decay is an antiviral defense in plants that is counteracted by viral RNA silencing suppressors. PLoS pathogens, 14(8), e1007228.

Li, F., & Wang, A. (2019). RNA-Targeted Antiviral Immunity: More Than Just RNA Silencing. Trends in microbiology, 27(9), 792–805.

Li, F., Zhao, N., Li, Z., Xu, X., Wang, Y., Yang, X., Liu, S. S., Wang, A., & Zhou, X. (2017). A calmodulin-like protein suppresses RNA silencing and promotes geminivirus infection by degrading SGS3 via the autophagy pathway in Nicotiana benthamiana. PLoS pathogens, 13(2), e1006213.

Li, P., Liu, C., Deng, W. H., Yao, D. M., Pan, L. L., Li, Y. Q., Liu, Y. Q., Liang, Y., Zhou, X. P., & Wang, X. W. (2019). Plant begomoviruses subvert ubiquitination to suppress plant defenses against insect vectors. PLoS pathogens, 15(2), e1007607.

Liu, L., Wang, H., Fu, Y., Tang, W., Zhao, P., Ren, Y., Liu, Z., Wu, K., & Zhang, X. (2023). Turnip crinkle virus-encoded suppressor of RNA silencing interacts with Arabidopsis SGS3 to enhance virus infection. Molecular plant pathology, 24(2), 154–166.

Liu, S., Wang, C., Liu, X., Navas-Castillo, J., Zang, L., Fan, Z., Zhu, X., & Zhou, T. (2021). Tomato chlorosis virus-encoded p22 suppresses auxin signalling to promote infection via interference with SKP1-Cullin-F-box^TIR1^ complex assembly. Plant, cell & environment, 44(9), 3155–3172.

Lukhovitskaya, N., & Ryabova, L. A. (2019). Cauliflower mosaic virus transactivator protein (TAV) can suppress nonsense-mediated decay by targeting VARICOSE, a scaffold protein of the decapping complex. Scientific reports, 9(1), 7042.

Mailliot, J., Vivoli-Vega, M., & Schaffitzel, C. (2022). No-nonsense: insights into the functional interplay of nonsense-mediated mRNA decay factors. The Biochemical journal, 479(9), 973–993.

May, J. P., & Simon, A. E. (2021). Targeting of viral RNAs by Upf1-mediated RNA decay pathways. Current opinion in virology, 47, 1–8.

May, J. P., Johnson, P. Z., Ilyas, M., Gao, F., & Simon, A. E. (2020). The Multifunctional Long-Distance Movement Protein of *Pea Enation Mosaic Virus 2* Protects Viral and Host Transcripts from Nonsense-Mediated Decay. mBio, 11(2), e00204–20.

May, J. P., Yuan, X., Sawicki, E., & Simon, A. E. (2018). RNA virus evasion of nonsense-mediated decay. PLoS pathogens, 14(11), e1007459.

Mei, Y., Ma, Z., Wang, Y., & Zhou, X. (2020). Geminivirus C4 antagonizes the HIR1-mediated hypersensitive response by inhibiting the HIR1 self-interaction and promoting degradation of the protein. The New phytologist, 225(3), 1311–1326.

Motomura, K., Le, Q. T., Hamada, T., Kutsuna, N., Mano, S., Nishimura, M., & Watanabe, Y. (2015). Diffuse decapping enzyme Dcp2 accumulates in Dcp1 foci under heat stress in Arabidopsis thaliana. Plant & cell physiology, 56(1), 107–115.

Niu, E., Ye, C., Zhao, W., Kondo, H., Wu, Y., Chen, J., Andika, I. B., & Sun, L. (2022). Coat protein of Chinese wheat mosaic virus upregulates and interacts with cytosolic glyceraldehyde-3-phosphate dehydrogenase, a negative regulator of plant autophagy, to promote virus infection. Journal of integrative plant biology, 64(8), 1631–1645.

Peng, J., Yang, J., Yan, F., Lu, Y., Jiang, S., Lin, L., Zheng, H., Chen, H., & Chen, J. (2011). Silencing of NbXrn4 facilitates the systemic infection of Tobacco mosaic virus in Nicotiana benthamiana. Virus research, 158(1-2), 268–270.

Perea-Resa, C., Hernández-Verdeja, T., López-Cobollo, R., del Mar Castellano, M., & Salinas, J. (2012). LSM proteins provide accurate splicing and decay of selected transcripts to ensure normal Arabidopsis development. The Plant cell, 24(12), 4930–4947.

Pérez-Vilaró, G., Scheller, N., Saludes, V., & Díez, J. (2012). Hepatitis C virus infection alters P-body composition but is independent of P-body granules. Journal of virology, 86(16), 8740–8749.

Popp, M. W., Cho, H., & Maquat, L. E. (2020). Viral subversion of nonsense-mediated mRNA decay. RNA (New York, N.Y.), 26(11), 1509–1518.

Qu, F., Ren, T., & Morris, T. J. (2003). The coat protein of turnip crinkle virus suppresses posttranscriptional gene silencing at an early initiation step. Journal of virology, 77(1), 511–522.

Qu, F., Ye, X., & Morris, T. J. (2008). Arabidopsis DRB4, AGO1, AGO7, and RDR6 participate in a DCL4-initiated antiviral RNA silencing pathway negatively regulated by DCL1. Proceedings of the National Academy of Sciences of the United States of America, 105(38), 14732–14737.

Rayson, S., Arciga-Reyes, L., Wootton, L., De Torres Zabala, M., Truman, W., Graham, N., Grant, M., & Davies, B. (2012). A role for nonsense-mediated mRNA decay in plants: pathogen responses are induced in Arabidopsis thaliana NMD mutants. PloS one, 7(2), e31917.

Seglen, P. O., & Gordon, P. B. (1982). 3-Methyladenine: specific inhibitor of autophagic/lysosomal protein degradation in isolated rat hepatocytes. Proceedings of the National Academy of Sciences of the United States of America, 79(6), 1889–1892.

Sun, H., Jing, X., Wang, C., Wang, P., Huang, Z., Sun, B., Li, P., Li, H., & Zhang, C. (2023). The Great Game between Plants and Viruses: A Focus on Protein Homeostasis. International journal of molecular sciences, 24(16), 12582.

Tong, X., Liu, S. Y., Zou, J. Z., Zhao, J. J., Zhu, F. F., Chai, L. X., Wang, Y., Han, C., & Wang, X. B. (2021). A small peptide inhibits siRNA amplification in plants by mediating autophagic degradation of SGS3/RDR6 bodies. The EMBO journal, 40(15), e108050.

Tsuzuki, M., Motomura, K., Kumakura, N., & Takeda, A. (2017). Interconnections between mRNA degradation and RDR-dependent siRNA production in mRNA turnover in plants. Journal of plant research, 130(2), 211–226.

Vidya, E., & Duchaine, T. F. (2022). Eukaryotic mRNA Decapping Activation. Frontiers in genetics, 13, 832547.

Wang, K., Fu, S., Wu, L., Wu, J., Wang, Y., Xu, Y., & Zhou, X. (2023). Rice stripe virus nonstructural protein 3 suppresses plant defence responses mediated by the MEL-SHMT1 module. Molecular plant pathology, 24(11), 1359–1369.

Wang, Y., Li, J., Wang, J., Han, P., Miao, S., Zheng, X., Han, M., Shen, X., Li, H., Wu, M., Hong, Y., & Liu, Y. (2022). Plant UVRAG interacts with ATG14 to regulate autophagosome maturation and geminivirus infection. The New phytologist, 236(4), 1358–1374.

Wu, J., Zhang, Y., Li, F., Zhang, X., Ye, J., Wei, T., Li, Z., Tao, X., Cui, F., Wang, X., Zhang, L., Yan, F., Li, S., Liu, Y., Li, D., Zhou, X., & Li, Y. (2023a). Plant Virology in the 21st Century in China: Recent Advances and Future Directions. Journal of integrative plant biology, 10.1111/jipb.13580.

Wu, K., Fu, Y., Ren, Y., Liu, L., Zhang, X., & Ruan, M. (2023b). Turnip crinkle virus-encoded suppressor of RNA silencing suppresses mRNA decay by interacting with Arabidopsis XRN4. The Plant journal: for cell and molecular biology, 116(3), 744–755.

Wu, K., Wu, Y., Zhang, C., Fu, Y., Liu, Z., & Zhang, X. (2021). Simultaneous silencing of two different Arabidopsis genes with a novel virus-induced gene silencing vector. Plant methods, 17(1), 6.

Xu, J., & Chua, N. H. (2009). Arabidopsis decapping 5 is required for mRNA decapping, P-body formation, and translational repression during postembryonic development. The Plant cell, 21(10), 3270–3279.

Xu, J., Yang, J. Y., Niu, Q. W., & Chua, N. H. (2006). Arabidopsis DCP2, DCP1, and VARICOSE form a decapping complex required for postembryonic development. The Plant cell, 18(12), 3386–3398.

Yang, M., & Liu, Y. (2022a). Autophagy in plant viral infection. FEBS letters, 596(17), 2152–2162. 10.1002/1873-3468.14349

Yang, M., Ismayil, A., Gao, T., Ye, Z., Yue, N., Wu, J., Zheng, X., Li, Y., Wang, Y., Hong, Y., & Liu, Y. (2023). Cotton leaf curl Multan virus C4 protein suppresses autophagy to facilitate viral infection. Plant physiology, 193(1), 708–720.

Yang, M., Ismayil, A., Jiang, Z., Wang, Y., Zheng, X., Yan, L., Hong, Y., Li, D., & Liu, Y. (2022b). A viral protein disrupts vacuolar acidification to facilitate virus infection in plants. The EMBO journal, 41(2), e108713.

Yang, M., Wang, Y., Li, D., & Liu, Y. (2022c). Plant virus infection disrupts vacuolar acidification and autophagic degradation for the effective infection. Autophagy, 18(3), 705–706.

Yang, M., Zhang, Y., Xie, X., Yue, N., Li, J., Wang, X. B., Han, C., Yu, J., Liu, Y., & Li, D. (2018). *Barley stripe mosaic virus* γb Protein Subverts Autophagy to Promote Viral Infection by Disrupting the ATG7-ATG8 Interaction. The Plant cell, 30(7), 1582–1595.

Zhang, X., & Guo, H. (2017). mRNA decay in plants: both quantity and quality matter. Current opinion in plant biology, 35, 138–144.

Zhang, X., Zhang, X., Singh, J., Li, D., & Qu, F. (2012). Temperature-dependent survival of Turnip crinkle virus-infected arabidopsis plants relies on an RNA silencing-based defense that requires dcl2, AGO2, and HEN1. Journal of virology, 86(12), 6847–6854.

Zhang, Z. J., Gao, Q., Fang, X. D., Ding, Z. H., Gao, D. M., Xu, W. Y., Cao, Q., Qiao, J. H., Yang, Y. Z., Han, C., Wang, Y., Yuan, X., Li, D., & Wang, X. B. (2020). CCR4, an RNA decay factor, is hijacked by a plant cytorhabdovirus phosphoprotein to facilitate virus replication. eLife, 9, e53753.

Zhang, Z., Chen, H., Huang, X., Xia, R., Zhao, Q., Lai, J., Teng, K., Li, Y., Liang, L., Du, Q., Zhou, X., Guo, H., & Xie, Q. (2011). BSCTV C2 attenuates the degradation of SAMDC1 to suppress DNA methylation-mediated gene silencing in Arabidopsis. The Plant cell, 23(1), 273–288.

